# Stitched peptides as potential cell permeable inhibitors of oncogenic DAXX protein

**DOI:** 10.1101/2022.09.25.508451

**Authors:** Clare Jelinska, Srinivasaraghavan Kannan, Yuri Frosi, Siti Radhiah Ramlan, Fernaldo Winnerdy, Rajamani Lakshminarayanan, Christopher J Brown, Anh-Tuan Phan, Daniela Rhodes, Chandra Verma

## Abstract

The death domain associated protein 6 (DAXX) is frequently upregulated in a number of common cancers where its suppression has been linked to reduced tumour progression. As a master regulator protein, with >70 reported protein interaction partners, the role of DAXX in its oncogenecity remains unclear. We designed and developed a set of novel stapled/stitched peptides that target a surface on the N-terminal helical bundle domain of DAXX which is the anchor-point for binding to multiple interaction partners (including Rassf1C, P53, Mdm2 and ATRX) and also for the auto regulation of the DAXX N-terminal SUMO interaction motif (SIM). We demonstrate that these peptides bind to and inhibit DAXX with an affinity higher than those reported for the known interaction partners and release the auto-inhibited SIM for interaction with SUMO-1. NanoBret assays show that the peptides enter cells and that their intracellular concentrations remain at nanomolar levels even after 24 hours, without causing membrane perturbation. Together our data suggest that these peptides are both tools for probing the molecular interactions of DAXX and potential precursors to the development of therapeutics.

## Introduction

The death domain associated protein 6 (DAXX) functions in several critical biological processes including cell death, cell survival, chromatin remodeling, transcription regulation, DNA repair and innate immunity.^1^ Increased DAXX expression has consistently been observed in diverse, epidemiologically prevalent cancers.^1^ How the participation of DAXX in a number of biological processes contributes to its oncogenicity is unclear. Moreover, whilst several *in vivo* studies in which DAXX overexpression is abrogated have demonstrated the potential of targeting DAXX in cancer therapy,^2–6^ to date, no such therapies exist. These observations highlight the importance of elucidating how DAXX orchestrates a complex network of molecular interactions to deepen our understanding of cancer biology, and evaluate the potential of DAXX as a therapeutic target.

First identified as a potentiator of FAS- induced apoptosis in the cytoplasm^7^, DAXX is found predominantly in the nucleus where it can be localized in a number of sub-cellular compartments such as promyelocytic leukaemia nuclear bodies (PML-NBs), the nucleoplasm, nucleoli and heterochromatin.^8–10^ The precise cellular role of DAXX appears to be dependent upon its ability to shuttle between compartments in a network of interactions often mediated by post-translational modifications, such as phosphorylation and SUMOylation.^11, 12^ For example, the role of DAXX in cell death has been linked to the phosphorylation dependent shuttling of DAXX between cytoplasm and nucleus, with pro- or anti- apoptotic outcomes depending upon increased fractionation of DAXX in the cytoplasm or nucleus, respectively. ^12, 13^ Interestingly, an increased cytoplasmic fractionation of DAXX has also been described as a favourable prognostic indicator in gastric cancer.^14, 15^ This is consistent with established cytoplasmic roles for DAXX in cell-death signalling pathways, but sub-fractionation of DAXX within the nucleus can also have important consequences for cell fate.^16^ For example, an increased association of DAXX with PML nuclear bodies relative to the nucleoplasm, a SUMOylation dependent process, has been reported to be pro-apoptotic and vice-versa.^16, 17^

Reversible SUMOylation is an important mechanism for the regulation of many proteins that interact with DAXX through dual SUMO interaction motifs (SIMs) located at its N- and/or C- termini (NSIM_DAXX_ and CSIM_DAXX_, respectively)^18^ and is, furthermore, an important regulatory mechanism for DAXX itself, which can also be SUMOylated. Specifically, whilst SUMOylated PML protein recruits DAXX to PML-NBs, DAXX must also be SUMOylated for full localisation to occur.^8, 19^ Functionally, the effect of sequestration of DAXX to PML-NBs appears to be a consequence of the specific factors involved and is thus context dependent. For example, sequestration of DAXX can either activate or repress transcription; DAXX, acting as a co-repressor has been shown to interact with multiple, functionally unrelated, SUMOylated transcription factors that are de-repressed upon increased association of DAXX with PML-NBs.^11, 19^ On the other-hand, as a transcriptional activator, sequestration of DAXX to PML-NBs can be repressive.^18^ The DAXX SIMs have been further implicated in recruitment to the nucleoli^10^ and other specific regions of the genome related to the histone chaperone activity of DAXX.^20^ Thus, the ability of DAXX to interact with multiple interaction partners, pathways and cellular compartments appears intimately related to reversible SUMOylation mechanisms.

DAXX can also affect transcription by acting directly on chromatin through its histone chaperone activity^21^ and/or the recruitment of epigenetic regulators such as HDACs^22^ and DNA methyltransferases.^23^ The ability to affect transcription through multiple pathways may in part explain the apparently diverse roles of DAXX, which has also been postulated as a potential cause of oncogenecity.^24^

An interesting nuclear pathway is the known interaction of DAXX with the chromatin remodeller ATRX, where DAXX functions as an H3.3 specific histone chaperone partner to ATRX.^25, 26^ Whilst the histone chaperone function of DAXX is conferred by the DAXX histone binding domain (DAXX_HBD_)^27, 28^ the anchor point for the interaction between ATRX and DAXX involves the DAXX-helical bundle domain (DHB_DAXX_).^29^ This domain of DAXX has also been reported to bind to multiple other protein interaction partners (Rassf1C^30^, Mdm2,^31^ HAUSP, p53 (and its homologues p63 and p73)^32^) in a mutually exclusive ‘partner-switch’ mechanism.^33, 34^ Importantly, interaction of NSIM_DAXX_ with SUMO is thought to be regulated allosterically by a self-interaction with a region on DHB_DAXX_ that also overlaps with the previously reported protein-protein interaction (PPI) surface.^35^ This strongly suggests that a shared DHB_DAXX_ PPI surface is central to the regulation of a complex network of interactions that modulate DAXX function. These observations taken together, suggest that generating specific inhibitors of the promiscuous DHB_DAXX_ PPI surface could be useful not only for probing the complex molecular assembly of DAXX with its partners *in vitro*, but also as potential DAXX-targeted therapeutics.

Atomic structures of DHB_DAXX_ in apo-, Rassf1C-bound and ATRX-bound forms show that the DHB_DAXX_ structure is comprised of the antiparallel packing of four alpha helices α1, α2, α4, and α5 and a short helix (α3) connecting helices α2 and α4.^33, 34, 36^ Although both Rassf1C and ATRX peptides bind between helices α2 and α5 of DHB_DAXX_, the helical axes of the bound peptides are drastically different, adopting near perpendicular orientations to each other in the two complexes. Significantly, the PPI surfaces occupied by the two peptides overlap with a hydrophobic pocket on the DHB_DAXX_ surface formed by residues V84, F87, Y124, V125 and I127. NMR chemical shift perturbation data also show significant changes in the amide cross-peak positions of these residues upon binding of DHB_DAXX_ to peptides derived from p53 and Mdm2 indicating that specific interactions are formed with the same hydrophobic surface in these complexes.^33^ Together, this strongly suggests that the hydrophobic pocket on the DHB_DAXX_ surface forms the anchor point for a shared PPI surface. For the purpose of targeting this pocket, which is quite shallow, a plausible strategy for generating inhibitors with high affinity and specificity is to identify peptidomimetic rather than small molecule inhibitors.^37^

In the study presented here aimed at discovering peptidomimetic PPI inhibitors of DHB_DAXX_ binding, we made use of the known interaction between ATRX and DHB_DAXX_ that has been reported to be stronger, by orders of magnitude, than that of other PPIs that have been characterized (p53, Mdm2, Rassf1C and NSIM_DAXX_).^33, 35^ The ATRX-derived peptide binds to DHB_DAXX_ in an extended alpha-helical conformation; a structure that lends itself well to a peptide stapling strategy.^38^ Based on this, we designed a series of such peptidomimetics using a combined analysis of biochemical, structural and molecular dynamics (MD) simulation data, and obtained a number of binders that bind with nano molar affinities. Our best stapled-peptide (based on experimentally determined thermodynamic binding parameters) binds to DHB_DAXX_ competitively with peptides derived from either ATRX or p53 and efficiently releases NSIM_DAXX_ from auto-inhibition so that it can bind to SUMO-1. These results validate the use of our peptidomimetic inhibitor/s to probe interactions between DHB_DAXX_ and multiple interaction partners *in vitro*. However, in cellular localisation studies, the stapled peptides were not able to enter the cytoplasm, becoming trapped in non-cytosolic endosomal compartments. To improve cell permeability, ‘stitched’ peptides^38^ were produced in which excess negative charges were also removed by mutation. These stitched peptides did enter cells and remained localized for >24 hours without outer membrane perturbation. What is less clear is whether the cytoplasmic concentration of the peptide, which is a few fold lower than the concentration required for binding of the peptides in vitro, are sufficient to bind to endogenous DAXX. In summary, our structural and biochemical evidence holds promise that ATRX-derived peptidomimetics targeting the ATRX-DAXX interface are effective inhibitors of multiple protein interactions that may contribute to the oncogenicity of DAXX.

## Results and Discussion

### Determination of the minimal DHB_DAXX_ binding sequence of ATRX, PEP_ATRX_

To enable the design of inhibitor peptides it was first necessary to define the minimal interaction surface between ATRX and DAXX. First, we dissected the previously reported DAXX Interacting Domain of ATRX (DID_ATRX_, residues 1190-1325)^29^ using a combination of experimental and computational techniques. We observed a robust interaction between DID_ATRX_ and the DAXX Helical Bundle (DHB_DAXX_, residues 55-144) using ITC (Kd 0.08 μM) (Figure S1 and Table 1). When the 136 amino acid long DID_ATRX_ sequence was used as query to retrieve structural templates from the protein databank database using BLASTp^39^ no template structure was identified. Instead, we used the automated I-TASSER pipeline^40^ to predict the atomic structure of DID_ATRX_. This method, which is well validated,^40^ is based on multi-threading alignment and iterative template fragment assembly simulation. It predicted the region of DID_ATRX_ spanning residues 1267 to 1284 to be helical, while the other parts of DID_ATRX_ were predicted to be disordered. This was consistent with the results from secondary structure and disorder predictions (Figure 1A).

**Figure 1.**
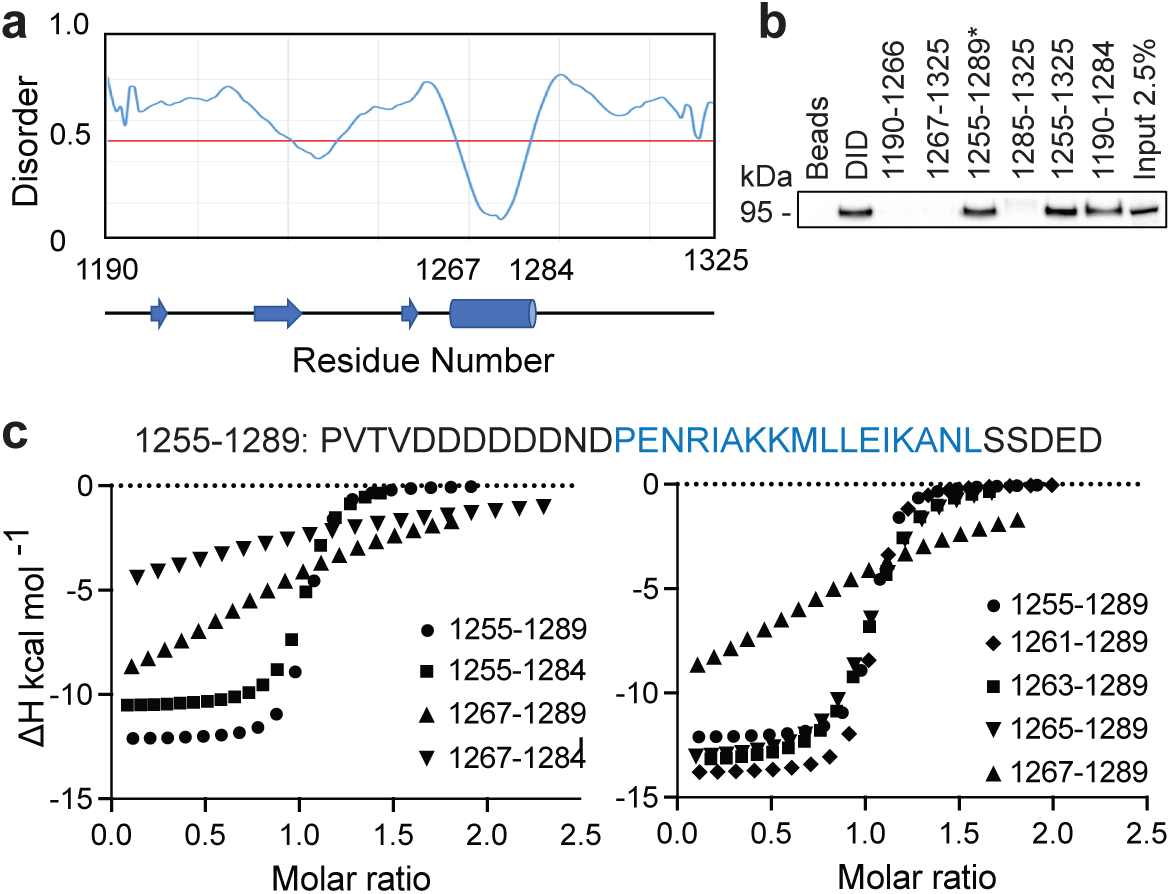
Dissection of DID_ATRX_ to define minimal sequence (PEP_ATRX_) required for specific binding to DHB_DAXX_. **A** Results of per-residue disorder prediction (disEMBL http://dis.embl.de) for DID_ATRX_ are plotted, with results from secondary structure prediction shown below. A helical region is predicted, spanning residues 1267-1284. **B** Western blot analysis of pull-down assays used to determine which fragments of DID_ATRX_ could pull down full-length DAXX protein from Hela nuclear extracts.‘*’ denotes the shortest fragment (1255-1289) found to bind to DAXX and used as the basis to test a series of further peptide fragments in ITC binding experiments. **C** The sequence of the 1255-1289 fragment is shown with the predicted helical region highlighted in blue. Truncations around the helical region were made to identify critical binding residues. Plots of binding enthalpies determined using ITC upon titration of these peptide fragments against DHB_DAXX_ are shown, where the identity of each peptide is indicated in the figure legend. Left panel: The results indicate that residues important for specific binding are contained within the N-terminal but not the C-terminal helix-flanking region. Right panel: The effect of systematically truncating the N-terminal flanking region, indicating that the two residues (N1265 and D1266) directly preceding the helix are critical for binding.

**Table 1.** Thermodynamic parameters from fitting ITC data obtained in DHB_DAXX_/ATRX peptide fragment binding experiments.

| Peptide | Kd (mM) | DH (kcal mol<sup>-1</sup>) | -TDS (kcal mol<sup>-1</sup>) | DG (kcal mol<sup>-1</sup>) |
| --- | --- | --- | --- | --- |
| 1255-1289 | 0.16 (0.01) | -12.2 (0.10) | 2.87 | -9.28 |
| 1255-1284 | 0.39 (0.03) | -10.6 (0.13) | 1.83 | -8.75 |
| 1261-1289 | 0.24 (0.02) | -13.8 (0.11) | 4.82 | -9.02 |
| 1263-1289 | 0.97 (0.09) | -13.4 (0.22) | 5.14 | -8.21 |
| 1265-1289 | 1.23 (0.08) | -13.3 (0.16) | 5.18 | -8.09 |
| 1267-1289 | 31.0 (7.64) | -11.9 (2.22) | 5.78 | -6.15 |
| 1267-1284 | 101 (162) | -9.3 (9.4) | 3.87 | -5.45 |
| 1264-1285 (PEP<sub>I</sub>) | 0.75 (0.07) | -13.3 (0.14) | 4.9 | -8.4 |
| DID<sub>ATRX</sub> (1190-1325) | 0.08 (0.01) | -11.7 (0.10) | 2.06 | -9.69 |

Calculated errors for fitted parameters are shown in parentheses.

The importance of the predicted helical region of ATRX in binding DAXX was confirmed experimentally in pull-down assays from Hela nuclear extracts. In this experiment a series of purified, recombinant GST-tagged DID_ATRX_ fragments were used to test for interaction with the full-length DAXX protein (Figure 1B). DAXX was detected in the pulldowns for all ‘helix’ containing fragments, with the exception of GST-ATRX (1267-1325). This result indicated that additional residues required for binding must be contained in the region preceding the alpha helix in DID_ATRX_. To further delineate the residues required for specific binding of DID_ATRX_ to DHB_DAXX_, we measured thermodynamic binding parameters for a series of DID_ATRX_ peptide fragments using ITC (Table 1). Plots of enthalpy change vs molar ratio were used to highlight regions and residues important for binding specificity (Figure 1C and D). The longest peptide in this series, ATRX (1255-1289) bound with an affinity similar to the full length DID_ATRX_ (Kds 0.16 μM and 0.08 μM, respectively). This peptide (Fig. 1B) includes 12 and 5 residues flanking the N- and C-terminal ends of the predicted alpha helix (1267-1284), respectively. We next assessed the contribution of these regions to binding (Figure 1C and Table 1). Whilst a negligible decrease in affinity (∼2-fold) was measured upon deletion of the C-terminal flanking region, loss of the N-terminal flanking region resulted in a dramatic reduction in binding (∼200 fold), a result consistent with the pull-down data (Fig. 1B). The N-terminal flanking sequence to the predicted alpha helix contains a polyaspartate sequence (1259-1266, punctuated by a single asparagine (N1265)), that we hypothesised to be important for binding to DHB_DAXX_ due to electrostatics and/or additional specific binding interactions. Systematic truncation of this region initially indicated modest losses of binding affinity (∼2-fold per DD removed). However, a ∼25-fold reduction in binding affinity, coupled with a dramatic shift in the enthalpy plot was observed upon deleting the two amino acid residues (N1265 and D1266) directly preceding the alpha helix. This indicated that one or both of these residues are necessary for binding specificity (Figure 1D and Table 1). Based upon these results we defined a peptide fragment of ATRX (1264- 1285), ‘PEP_ATRX_’, sufficient for a specific interaction with the DHB_DAXX_. The measured binding constant for the interaction between PEP_ATRX_ and DHB_DAXX_ (Kd 0.75 μM) is ∼ 10- fold weaker than that for the corresponding interaction with the longer DID_ATRX_ (Kd 0.08 μM) (Table 1).

### Modelling the structure of the PEP_ATRX_ – DHB_DAXX_ complex

The only available atomic structures when we began our inhibitor design studies were the solution NMR structures of apo-DHB_DAXX_ and DHB_DAXX_ in complex with a peptide derived from the Rassf1C (PEP_Rassf1C_) tumour suppressor.^33^ PEP_Rassf1C_ adopts a helical conformation upon binding to DHB_DAXX_ and given this, together with our analyses (above) suggested that PEP_ATRX_ also binds to DHB_DAXX_ as an alpha helix.

A helical model of PEP_ATRX_ was generated (using I-TASSER, as described above) and docked into the Rassf1C binding pocket of DHB_DAXX_ using the program HADDOCK (High Ambiguity Driven biomolecular DOCKing).^41, 42^ The docking protocol was established by docking the Rassf1C peptide into the binding pocket of DHB_DAXX_ to successfully reproduce the bound conformation of the peptide from the NMR structure with rmsd between the top docking solution and the 1^st^ conformation of the NMR ensemble (2KZU.PDB) of ∼1.5 Å. Additional docking runs of the PEP_ATRX_ helix (obtained from denovo modeling) against DHB_DAXX_ were then performed. Several docked poses of the peptide were generated, yielding 7 major clusters of the peptide bound to the same pocket of DHB_DAXX_ (Figure S2A). To further refine the bound poses and remove the less stable binders, a representative complex from each cluster was subjected to molecular dynamics (MD) simulations at a temperature of 330K. When the NMR structure of DHB_DAXX_/PEP_Rassf1C_ was subjected to this protocol, the peptide was stably bound (rmsd ∼ 3 Å against the starting conformation), with no peptide unbinding/protein unfolding events witnessed during the simulations (Figure S2B). In contrast, only 2 of the 7 docked models of DHB_DAXX_/PEP_ATRX_ remained stably bound and the conformations of these two remaining poses were found to overlap (Figure S2C). Remarkably, the positioning of these PEP_ATRX_ poses on the DHB_DAXX_ surface differed substantially from the DHB_DAXX_ surface occupied by PEP_Rassf1C_ in the solution structure with the helical axes juxtaposed by ∼90° (Figure 2A).

**Figure 2.**
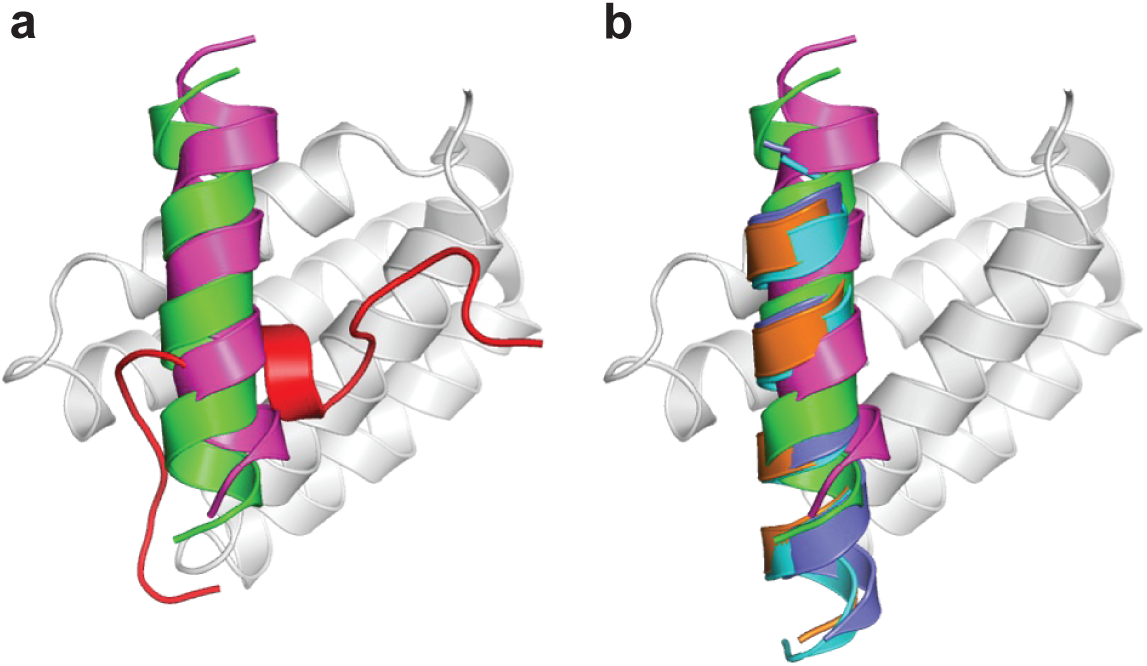
Comparison of the modelled DHB_DAXX_/PEP_ATRX_ structure with experimentally determined DHB_DAXX_/PEP_Rassf1C_ and DHB_DAXX_/PEP_ATRX_ structures. Cartoon representations comparing the 2 docked poses of PEP_ATRX_ (green and magenta) in the modeled DHB_DAXX_/PEP_ATRX_ structures with **A,** the solution structure of DHB_DAXX_/PEP_Rassf1C_(red) (pdb 2KZU) and **B,** the co-crystal structures of DHB_DAXX_/PEP_ATRX_ (pdb 5Y18 (orange), 5GRQ (blue) and 5Y60 (cyan)). The structures were superimposed using the bound DHB_DAXX_ (grey) backbone.

During the course of this study three X-ray structures of DHB_DAXX_ in complex with various, overlapping ATRX peptides were published.^34, 36, 43^ In all these structures, residues 1267- 1283 of ATRX were found to adopt a helical conformation. In two crystal structures that contained a flanking region preceding the helix, H-bonds were observed between the side- chains of D1266_ATRX_ and K122_DAXX_, also explaining our observations indicating that residues preceding the helix are important for specific binding. Remarkably, the two stably docked poses from our in-silico modelling aligned well with the bound ATRX peptide in the crystal structures (rmsd ∼ 2 Å) despite the juxtaposed orientation of PEP_ATRX_ in the initial model, demonstrating the robustness of our de-novo modelling approach (Figure 2B). For further PPI inhibitor design studies, we made use of the structural information from the crystal structure of the DHB_DAXX_/PEP_ATRX_ complex (PDB, 5GRQ), because it provides more comprehensive information on the interaction than our modelling. From this point we renamed the PEP_ATRX_ template sequence as ‘PEP_I_’ and renumbered the residues according to the peptide sequence (1-22). Details of the specific interactions formed between PEP_I_ and DHB_DAXX_ together with the surface charge complementarity are shown in Figure 3.

**Figure 3.**
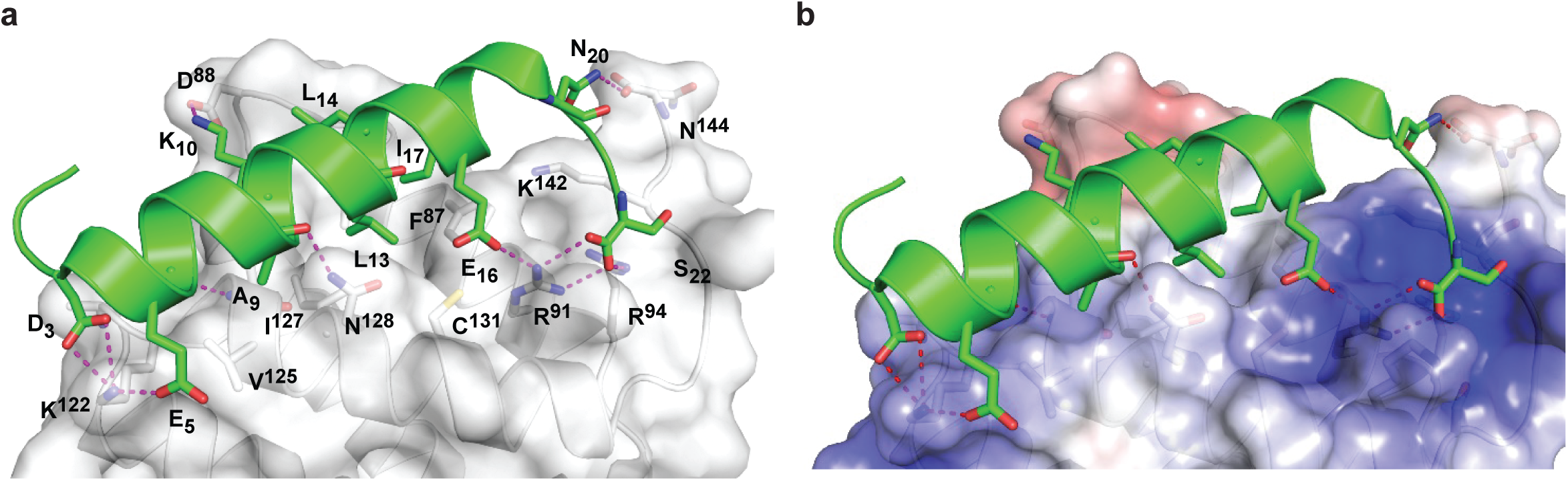
Structural views of DHB_DAXX_ – PEP_ATRX_ interactions. A. Cartoon representation of PEP_ATRX_ (green) bound to DHB_DAXX_ surface (grey/white). The interacting protein-peptide residues are labelled and highlighted as sticks. **B** As in **A** but with labels removed and displaying the electrostatic surface potential of the DHB_DAXX_ protein (red to blue colours indicating a range from -5 to +5 kcal/mol). This highlights the charge complementary of the interaction surface to PEP_ATRX_ residues and locates the hydrophobic pocket on DHB_DAXX_ that is important for binding both PEP_Rassf1C_ and PEP_ATRX_.

### Conformational dynamics of PEP_I_ and DHB_DAXX_ in apo and bound states

To design tight binding analogues of PEP_I_ it is necessary to understand the behaviour of the peptide and target alone in solution, as it provides a picture of the thermodynamic parameters that could be tuned towards designing higher affinity analogues. Thus, the conformational behaviour of unbound PEP_I_ in solution was explored using Hamiltonian Replica Exchange Molecular Dynamics (HREMD). In simulations, initiated using the crystallographic helical structure, PEP_I_ rapidly (< ∼5 ns) lost its secondary structure and became highly flexible (Figure S3A), with a conformational landscape populated by largely disordered states characterized by transient helical motifs (∼15% helicity overall). In comparison, the experimentally determined helical propensity, measured using Circular Dichroism spectropolarimetry (CD) was far lower (∼1.5%) in solution (Figure S3B and Table 2). This underscores that binding of PEP_I_ to DHB_DAXX_ in an alpha helical conformation is driven by strong enthalpic forces.

**Table 2.** Thermodynamic parameters from ITC experiments for the binding of SPEP1-7 to DHB_DAXX_. Additionally, percentage helicity of isolated peptides, calculated from CD spectra are included.

| Peptide | K<sub>d</sub> (mM) | DH (kcal mol<sup>-1</sup>) | -TDS (kcal mol<sup>-1</sup>) | DG (kcal mol<sup>-1</sup>) | Helicity (%) |
| --- | --- | --- | --- | --- | --- |
| PEP<sub>I</sub> | 0.75 (0.07) | -13.3 (0.14) | 4.94 | -8.36 | 1.4 |
| SPEP1 | 0.09 (0.01) | -9.0 (0.17) | -0.57 | -9.63 | 17.4 |
| SPEP2 | 0.05 (0.01) | -10.2 (0.12) | 0.3 | -9.9 | 11.7 |
| SPEP3 | 0.18 (0.04) | -8.8 (0.20) | -0.39 | -9.19 | 17.9 |
| SPEP4 | 0.34 (0.05) | -7.6 (0.17) | -1.3 | -8.9 | 13.9 |
| SPEP5 | 0.55 (0.08) | -7.4 (0.25) | -1.1 | -8.5 | 8.8 |
| SPEP6 | 0.07 (0.01) | -8.5 (0.18) | -1.2 | -9.8 | 14.2 |
| SPEP7 | 0.08 (0.07) | -12.8 (0.11) | 3.1 | -9.7 | 23.4 |

**Calculated errors for fitted parameters are shown in parentheses.**

The structural dynamics of apo DHB_DAXX_ in solution were explored using classical MD simulations and, except for the N- and C- termini residues (55-60 and 137-144, respectively), was found to remain stable (rmsd < 2 Å(Figure S3C). This is in accord with the solution structure of apo DHB_DAXX_ (2KZS.PDB).

MD simulations of the DHB_DAXX_/PEP_I_ complex showed that the conformations of both DHB_DAXX_ and PEP_I_ remained stable (RMSD ∼2.5 Å (Figure S3C)). The flexibility of the C-terminal region of apo DHB_DAXX_ was attenuated in the complex by formation of salt-bridge/hydrogen bond interactions with PEP_I_. Flexibility at the N-terminal region of DHB_DAXX_ upon complexation was unaffected. Whilst bound, PEP_I_ largely retained its α-helical conformation during the simulation (∼70% α-helicity), with high flexibility observed towards the C- terminal region of the peptide, which remained disordered throughout (Figure S3C). The hydrophobic and charged interactions observed in the co-crystal structure of DHB_DAXX_/PEP_I_ (Figure 3) were retained for ∼80% of the MD simulation. Together these results indicate that binding of PEP_I_ to DHB_DAXX_ is driven by the strong enthalpic forces and hence complex-formation requires the folding of the PEP_I_ peptide.

### Design of stapled peptide inhibitors of DHB_DAXX_

The low helicity of PEP_I_ alone in solution shows that PEP_I_ binds to DHB_DAXX_ as an induced alpha helix. Thus, constraining unbound PEP_I_ into an alpha helical conformation resembling its bound state should increase the binding affinity to DHB_DAXX_ by reducing the entropic cost of binding.^44^ To achieve this, we sought to design stapled peptide analogues of PEP_I_. To identify appropriate positions on the PEP_I_ peptide for the introduction of the hydrocarbon linkers, we first determined the residues in the peptide which, upon mutation, would result in minimal perturbation of the PEP_I_/DHB_DAXX_ interaction.

MD simulations of the DHB_DAXX_/PEP_I_ complex were mined to decompose the contributions of each residue of PEP_I_ to the total binding energy (Figure 4A). As expected, the charged/polar residues (D3, E5, N6, K10, E16, and S22), and hydrophobic residues (A9, L13, L14, I17, and L21) at the DHB_DAXX_/ PEP_I_ interface seen in the crystal structure of the complex (Fig 3) contribute favourably to binding.

**Figure 4.**
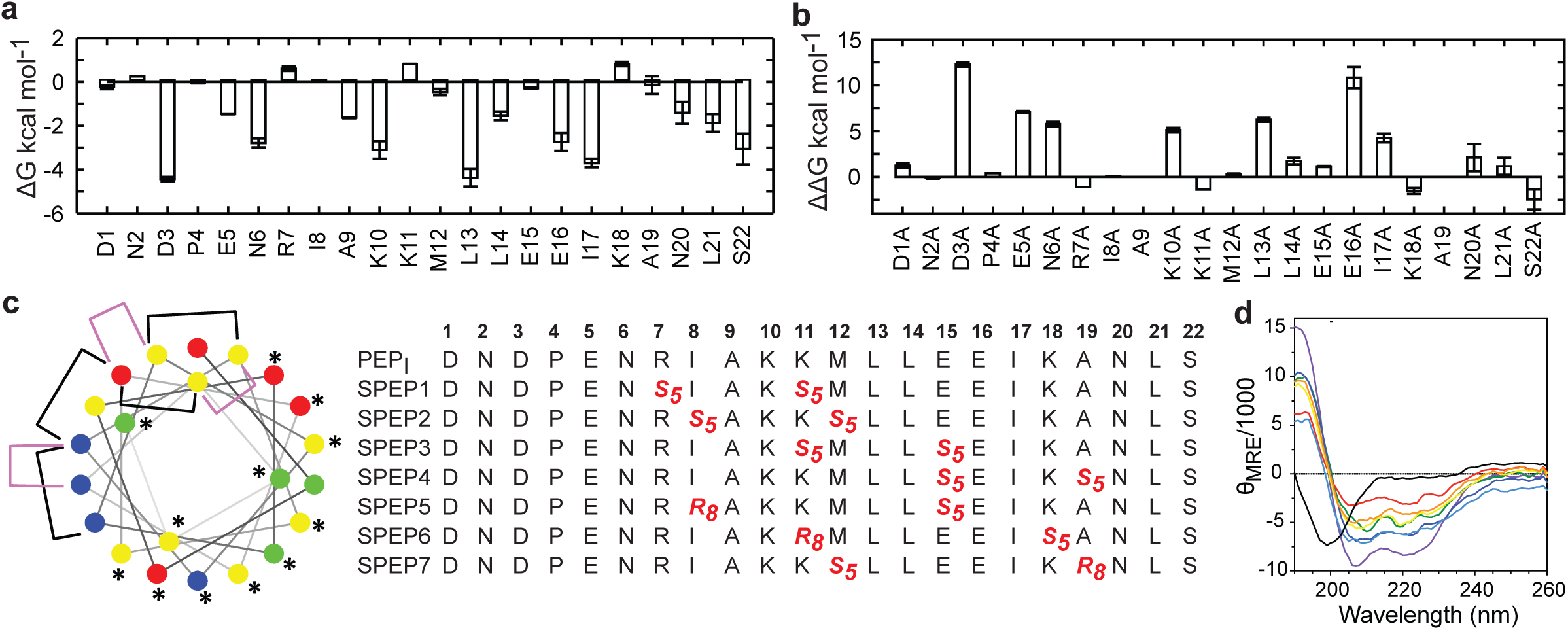
In-silico analysis of DHB_DAXX_–PEP_I_ key interacting residues and stapled peptide designs. A. Per-residue binding free energy contributions of PEP_I_ in the DHB_DAXX_/PEP_I_ complex from MD simulations. **B** Computational alanine scanning of PEP_I_ residues in the DHB_DAXX_/PEP_ATRX_ complex. **C** Helical wheel representation of the linear PEP_ATRX_ peptide sequence used for the design of stapled peptides. Colour-coded circles indicate polar/acidic (red), polar/basic (blue), polar/uncharged (green) and nonpolar (yellow) residues. Residues important for interaction with DHB_DAXX_ are highlighted with asterisks. Positions bridged by hydrocarbon linkers i,i+4 or i,i+7 are indicated with magenta or black brackets, respectively. Sequences of stapled peptides (SPEP1-7) based on MD simulations are shown alongside using standard nomenclature with staple positions highlighted in red. Residue number is shown in bold above. **D** CD spectra of all stapled peptides (various colours) demonstrate a marked increase in helicity compared to the linear PEP_I_ peptide (black).

The contribution of each amino acid residue in PEP_I_ to binding was also explored by carrying out *in-silico* alanine scanning substitutions. These results (Fig. 4B) mirrored the per-residue contributions obtained from the decomposition analysis, where major loss in affinity (corresponding to ΔΔG > 2 kcal/mol) was observed for alanine mutations at residues D3, E5, N6, K10, E16, L13 and I17. In contrast, the other positions were quite tolerant to alanine substitution, with substitution at some positively charged residues (R1270, Lys 1274 and Lys1281) even increasing binding affinity. Overall, these studies suggested several positions where staples could be incorporated without significant perturbations to target engagement.

The incorporation of staples also requires careful selection of sidechains with appropriate stereochemistry and we used standard strategies that have proven effective for stapling right-handed alpha-helices (*i.e.,* S5 to S5 for (i, i+4) linkages, and R8 to S5 for (i, i+7) linkages).^45^ Using these linkages and the simulations to guide staple placement, we designed 7 stapled peptides, referred to as SPEP 1-7 (Figure 4C).

### Peptide stapling increases helicity

To validate the design of our stapled peptides, HREMD simulations were carried out on SPEP 1-7 peptides in solution, starting from alpha helical conformations. As expected, all stapled peptides were predicted to have greater overall helicities in solution, ranging from 26-58%, compared to the linear PEP_I_ peptide (∼15%) (Fig S4). Analysis of per-residue helicity also suggested that the stapled peptides tend to retain helicity in the centres of the peptides with increased flexibility at the termini (Fig S4). No correlation was apparent between changes in helicity and the position of the staples, as has also been reported in studies of other systems.^46, 47^

The helicities of the linear PEP_I_ and SPEP1-7 peptides were determined experimentally using CD spectropolarimetry (Fig. 4D and S4). Whilst confirming a substantial increase in helicity upon stapling (ranging from 9-23%) compared to the linear PEP_I_ peptide (1.5%), the experimental values obtained were consistently lower than values obtained from simulations (Fig S4).

### Modelling of DHB_DAXX_ - SPEP complexes

The binding of SPEP 1-7 peptides to DHB_DAXX_ was modelled using the co-crystal structure of the DHB_DAXX_/PEP_ATRX_ complex [PDB 5GRQ, ATRX residues 1264-1285 (PEP_I_), Fig. 3] as a starting point. Staples were individually modelled into the PEP_I_ peptide and then subjected to MD simulations for 100 ns (each in triplicate). The SPEP 1-7 peptides all remained stable during the MD simulations with rmsd values reaching ∼2 Å from the corresponding starting conformation (Fig. S5). The introduction of hydrocarbon linkers increased the helicity of the bound peptides, which retained 75 to 85% helicity throughout the simulations. Flexibility was observed at both termini of the peptides even in their bound states, especially at the C-terminal ends which generally showed less helicity compared to the rest of the peptide; exceptions to this were SPEP6 and SPEP7 where the staple linker spans the C- terminal region of PEP_I_. The bound conformations of DHB_DAXX_ were also stable with rmsd ∼3 Å from the corresponding starting conformation (Fig. S5).

Structural representations of MD snapshots of the complexes (Fig. 5) show that the hydrocarbon staples remained exposed to solvent without engaging the DHB_DAXX_ surface. In accordance with the crystal structure of DHB_DAXX_/PEP_I_ (Fig. 3), the hydrophobic residues of SPEP 1-7 including A9, L13, and I7 remained buried with L14 only partially buried in the hydrophobic pocket/binding site of DHB_DAXX_. In contrast, L14 is exposed to solvent in simulations of the unstapled DHB_DAXX_/PEP_I_ complex (not shown). Salt bridges and hydrogen bond interactions observed in the crystal structure of the DHB_DAXX_/PEP_I_ complex were also well retained (∼90%) throughout the simulations of the DHB_DAXX_/SPEP complexes. In addition, several new interactions were observed during MD simulations of the DHB_DAXX_- SPEP complexes (since they have not been verified experimentally we do not detail them here). Overall, charged and polar residues D3, E5, N6, K10, E16, S22 along with the hydrophobic residues A9, L13, L14, I17, L21, contributed the most to the binding of the stapled peptides in the MD simulations (Fig. S6). These analyses allowed us to identify a number of promising stapled peptides with satisfactory overall binding parameters (Table S1).

**Figure 5:**
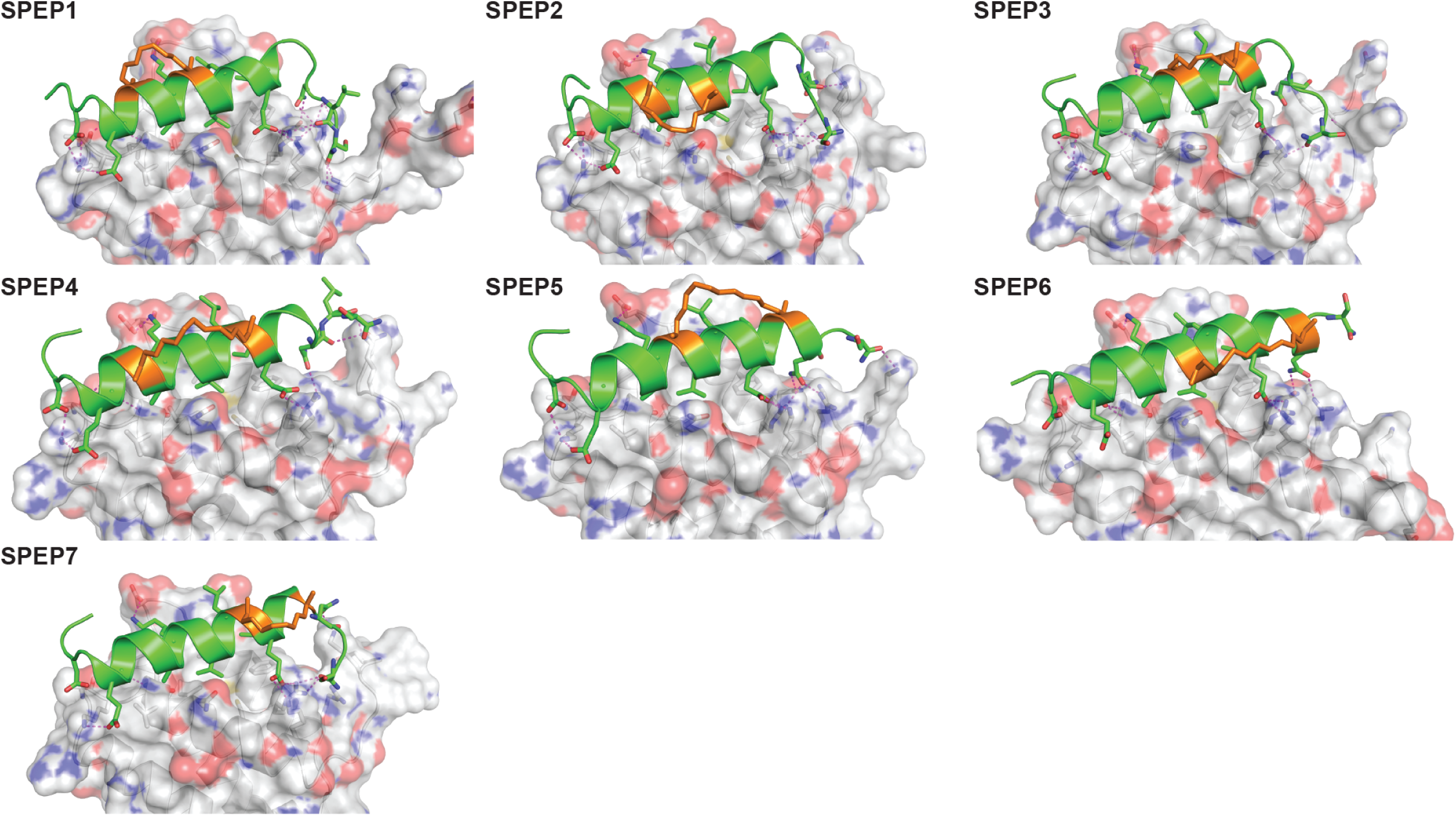
Structural representations of MD snapshots of DHB_DAXX_/SPEP complexes. Structural models for DHB_DAXX_ complexes with **A** SPEP1, **B** SPEP2, **C** SPEP3, **D** SPEP4, **E** SPEP5, **F** SPEP6 and **G** SPEP7 are shown. The DHB_DAXX_ (white) and bound peptide (green) conformations are shown as surface and cartoon representations, respectively with interacting residues and staples (orange) highlighted in sticks. Hydrogen bonds are indicated by dotted lines (magenta).

These MD analyses predict the formation of stable complexes of each of the SPEP peptides with DHB_DAXX_ in which the interface interactions closely resemble those formed between DHB_DAXX_ and the linear PEP_I_. Accordingly, all 7 SPEP designs were synthesised for experimental determination of binding affinity.

### Peptide stapling increases binding affinity to DHB_DAXX_

Having identified promising stapled peptides from the modelling, the binding affinities of the stapled peptides to DHB_DAXX_ were analysed using ITC. These measurements showed that SPEP 1-7 bound to DHB_DAXX_ with Kd values in the ∼ 50 nM – ∼1 μM range (Table 2, Figure S7). All seven stapled peptides bound to DHB_DAXX_ with similar or higher affinity compared to the linear PEP_I_ with four of the peptides, SPEP 1, 2, 6 and 7, binding ∼10-20 fold more strongly. Although the stapled peptides bound to DHB_DAXX_ with free energy gains of up to 1.5 kcal/mol, the linear PEP_I_ bound with more favourable enthalpic contributions (Table 2). The linear peptide is not structured in solution and favourable enthalpy compensates for the entropic penalty paid when the disordered peptide adopts a helical conformation upon binding to DHB_DAXX_. The lack of structure and associated flexibility enables the peptide to maximise potential interactions with the target, thus resulting in high enthalpy. In comparison, the stapled peptides are conformationally trapped in alpha helical conformations and binding to DHB_DAXX_ is expected to incur smaller entropic penalties. The reduction in internal flexibility of the stapled peptides could however result in fewer interactions with the target resulting in lower associated enthalpies. However, the computed distributions of the contacts between the peptides and the target did not yield a clear pattern. Since most interactions are mediated by exposed sidechains, this is not surprising; there will be the associated complexity of solvent interactions which are difficult to quantify precisely. Indeed, positive entropic contributions to binding were measured experimentally for 5 of the 7 peptides suggesting unexpected increases in disorder upon binding. While we did not see a correlation between the flexibility of the peptides in the bound state simulations and the experimentally measured order of entropies ie 6>4>5>1>3, our MD snapshots of the complexes suggests that the peptides with higher entropy all have staples pointing into the solvent (Fig. 5). It is quite possible that this exposure of the hydrophobic region into a polar solvent makes them very frustrated, resulting in higher entropies. In addition there is also the likelihood of differential release of water molecules as they are displaced from the binding pocket of DHB_DAXX_. Such increases in binding entropy could also indicate a loss of binding specificity and indeed, this behaviour was always accompanied by losses in binding enthalpy. Since the preservation of binding specificity is the most important factor in the development of our inhibitors, we ranked the four best binders wrt binding affinity (Kd) according to the binding enthalpy (ΔH) as follows: SPEP7>SPEP2>SPEP1>SPEP6.

### Binding competition experiments

To test whether our stapled peptides could inhibit multiple protein interaction partners that bind to DHB_DAXX_, thus providing a means to disrupt DAXX function, we performed competition experiments using a fluorescence polarization assay. First, using an N-terminally FAM-modified version of SPEP7 (FAM-SPEP7), we determined that this peptide binds to DHB_DAXX_ with a comparable affinity to the non-fluorescent SPEP7 (Kd 46 nM vs 80 nM, respectively) (Fig. 6A). This result demonstrated that the FAM-label doesn’t interfere with binding and validated use of the fluorescence polarization assay in subsequent experiments. FAM-SPEP7 could also be displaced from DHB_DAXX_ by the non-fluorescent SPEP1-7 peptides with comparable affinities to those measured by ITC confirming that these peptides are all targeting the same interaction site (Fig. S8). These results indicate that FAM-SPEP7 is a good model peptide for probing interactions of DHB_DAXX_ with other interaction partners (Fig. 6). By titrating DID_ATRX_ or p53 peptide competitors against the DHB_DAXX_/FAM-SPEP7 complex, we confirmed that the FAM-SPEP7 peptide could be displaced by either ligand (Fig. 6B). A similar binding affinity for the interaction between DID_ATRX_ and DHB_DAXX_ was obtained from the competition data compared to direct binding measured using ITC (Kd 193 nM vs 80 nM, respectively), suggesting high complementarity between the interaction surfaces, as expected. However, we estimate that competitive binding of the p53 peptide to DHB_DAXX_ is ∼10 fold weaker in competition experiments compared to direct binding. This disparity is likely due to a low complementarity of the SPEP7 and p53 peptide interaction surfaces on DHB_DAXX_, where the overlap between binding sites may be largely limited to the hydrophobic pocket that characterises the shared PPI interface.^36^ In conclusion, our stapled peptides compete well with multiple DAXX interaction partners *in vitro* and hence have the potential to disrupt DAXX interactions *in vivo*. In addition, protein interaction partners that bind to DHB_DAXX_ with low complementarity compared to the SPEP7/DHB_DAXX_ interaction will be inhibited disproportionately strongly.

**Figure 6.**
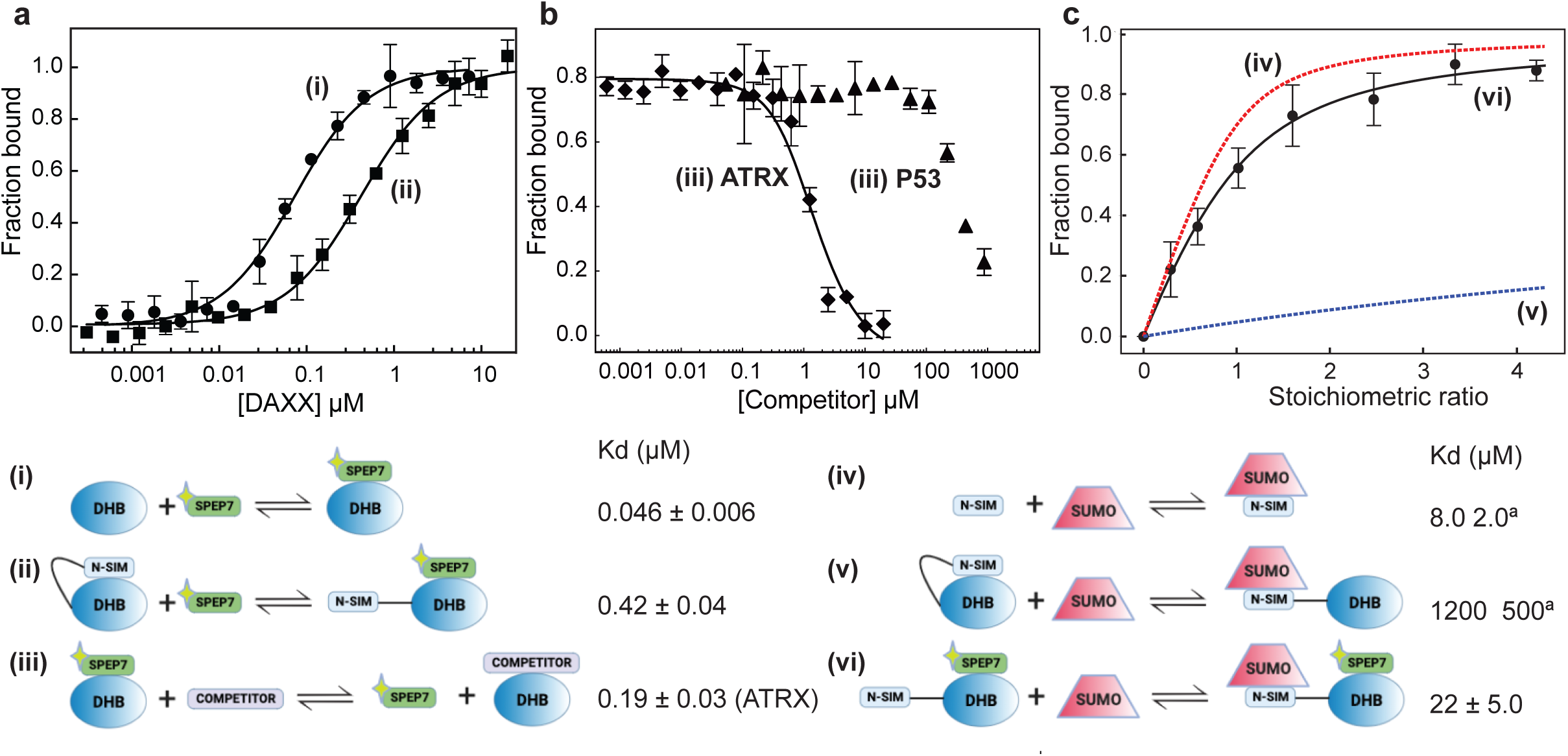
Multiple interactions and binding modes of DHB_DAXX_ probed using FAM- SPEP7. Data depicted in **A-C** are labeled according to the relevant schematic ((i) to (vi)) that describes the corresponding binding interaction and associated Kd. A yellow star is used to indicate the FAM moiety. **A** Saturation binding of FAM-SPEP7 to DHB_DAXX_ ((i), closed circles) or NSIM-DHB_DAXX_ ((ii), closed squares). **B** (iii) Competitive displacement of FAM- SPEP7 from DHB_DAXX_ by either DID_ATRX_ (closed diamonds) or PEP_P53_ (closed triangles). The data points in **A** and **B,** shown with standard error bars, are averages of three independent measurements. **C** Release of NSIM_DAXX_ for binding to SUMO by FAM-SPEP7 ((vi) closed circles)). The data points with standard error bars are the normalized average chemical shift change observed for 7 cross-peaks in the 1H-15N HSQC spectrum of the ^15^N- NSIM-DHB_DAXX_/FAM-SPEP7 complex upon titration with SUMO1. An apparent Kd for the interaction was obtained by fitting the curve. Simulated binding curves (dashed lines) for NSIM_DAXX_ /SUMO1 ((iv) red) and NSIM-DHB_DAXX_/SUMO1 ((v) blue dashed) interactions were calculated from published data ^35^.

### Release of NSIM_DAXX_ auto-inhibition

It has been proposed that the N-terminal SUMO Interaction Motif of DAXX (NSIM_DAXX_) can be sequestered from interaction with SUMO by self-interaction with DHB_DAXX_ *in vitro*.^35^ This is likely to be an important regulatory mechanism that prevents promiscuous interactions between NSIM_DAXX_ and non-cognate SUMOylated protein interaction partners. Release of the NSIM_DAXX_ could provide a mechanism by which DAXX could be sequestered (e.g to PML bodies) and thereby provide a general means of inhibition in cells that aberrantly upregulate DAXX.^11, 18^ We therefore tested the hypothesis that binding of our stapled peptide inhibitors could be used to release NSIM_DAXX_ for interaction with SUMO-1 (NSIM_DAXX_ has no binding preference for a particular homolog of SUMO ^35^), using SPEP7 as a model. Saturation binding of FAM- SPEP7 to NSIM-DHB_DAXX_ was quantified using fluorescence polarisation (Fig. 6A). Consistent with auto-regulation, we found that the binding affinity was ∼10 fold weaker when compared to binding to DHB_DAXX_ alone (416 nM vs 45 nM, respectively) (Fig. 6A). To demonstrate that NSIM_DAXX_ is released from DHB_DAXX_ upon binding of SPEP7, we then performed a series of titrations, monitored using ^1^H-^15^N HSQC NMR spectroscopy to detect specific interactions. We first titrated ^15^N-labelled NSIM-DHB_DAXX_ with FAM-SPEP7 and observed discrete changes in amide cross-peaks, consistent with the formation of a specific complex [Fig. S9A]. Upon titration of the pre-formed, saturated, ^15^N-NSIM-DHB_DAXX_/FAM- SPEP7 complex with unlabelled SUMO-1, we observed further specific chemical shift perturbations in either intermediate or fast exchange. A total of 9 amide cross-peaks in fast exchange were used to determine the binding affinity of SUMO-1 to NSIM_DAXX_ in context of the NSIM-DHB_DAXX_/FAM-SPEP7 complex (Fig. S9B). Fitting of the normalised, averaged data from these 9 cross-peaks yielded an apparent Kd of ∼22 μM (Fig. 6C). This is comparable to the affinity reported for direct interaction between SUMO-1 and the isolated NSIM_DAXX_ peptide (8 μM) and is ∼50 fold stronger than the interaction reported for binding of SUMO-1 to NSIM-DHB_DAXX_ in the absence of a DHB_DAXX_ binding inhibitor (1.2 mM).^35^ This clearly demonstrates that NSIM_DAXX_ can be released for binding to SUMO-1 (i.e SUMOylated interaction partners) through displacement from the DHB_DAXX_ by the stapled peptide SPEP7.

### Strategy for enhancing cell permeability of SPEP Peptides

Our *in vitro* experiments showing that stapled peptides, SPEP 1, 2, 6 and 7 bind tightly to DHB_DAXX_ and that FAM- SPEP7 (as an example) competes with peptide ligands derived from cognate interaction partners and can also release NSIM_DAXX_ from auto-regulation were encouraging. Therefore, we went on to test the cell permeability of our stapled peptides. We detected no cytotoxicity as measured in LDH release assays when HCT116 cells were treated with up to 50 uM SPEP1-7 (Fig. S10A). However, live cell imaging experiments using FAM- SPEP7, showed that the peptide was retained in non-cytosolic compartments suggesting that this peptide would not be able to reach and target endogenous DAXX (Fig. S10B). To improve entry of the SPEP peptides into the cytoplasm and facilitate target engagement with DAXX, the following physiochemical properties, that have been shown to be critically important for cellular permeability, were explored: Charge, hydrophobicity and amphipathicity.^47, 48^ The SPEP1-7 peptides are derived from a highly charged PEP_I_ template sequence, containing an abundance of aspartate, glutamate and lysine residues and carrying a net negative charge overall. The energetic costs of desolvating charged residues in order to traverse the cells lipid membrane bilayer can often be refractory to cellular uptake.^49^ Furthermore, a net positive charge is generally considered favourable for the engagement of negatively charged cell membranes. Therefore, we decided to test whether substitutions and/or deletion of charged residues not involved in binding would improve permeability. We simultaneously investigated whether increasing the hydrophobic and amphipathic character of our stapled peptides by incorporation of an additional staple could confer improvements in binding, stability and cellular activity. Such double-stapled peptides, designed with a common attachment/anchoring point resulting in contiguous hydrocarbon staples are referred to as “stitched” peptides and recent work has reported improvements in thermal and chemical stability, proteolytic stability, increased helicity and cell permeability of stitched peptides.^47, 50^

### Design of stitched peptides (STPEP) for improved cell permeability

The binding data obtained for SPEP1-7 peptides and DHB_DAXX_ suggested obvious pairs of staples that could be used to design two (i, i+4; i+4, i+7) stitched configurations by combining either SPEP1+6 or SPEP2+7, where the position of the i, i+4 staples in SPEPs 1 and 2 match the sites used for the i, i+7 staples in SPEPs 6 and 7, respectively. With additional deletions and mutations included to remove negative charges, a total of seven stitched peptides STPEP 1-7 were designed (Fig. 7). To efficiently utilise peptide synthetic bandwidth, computational modelling of the stitched peptides was then used to predict helicity of the isolated peptides and their binding to DHB_DAXX_.

**Figure 7.**
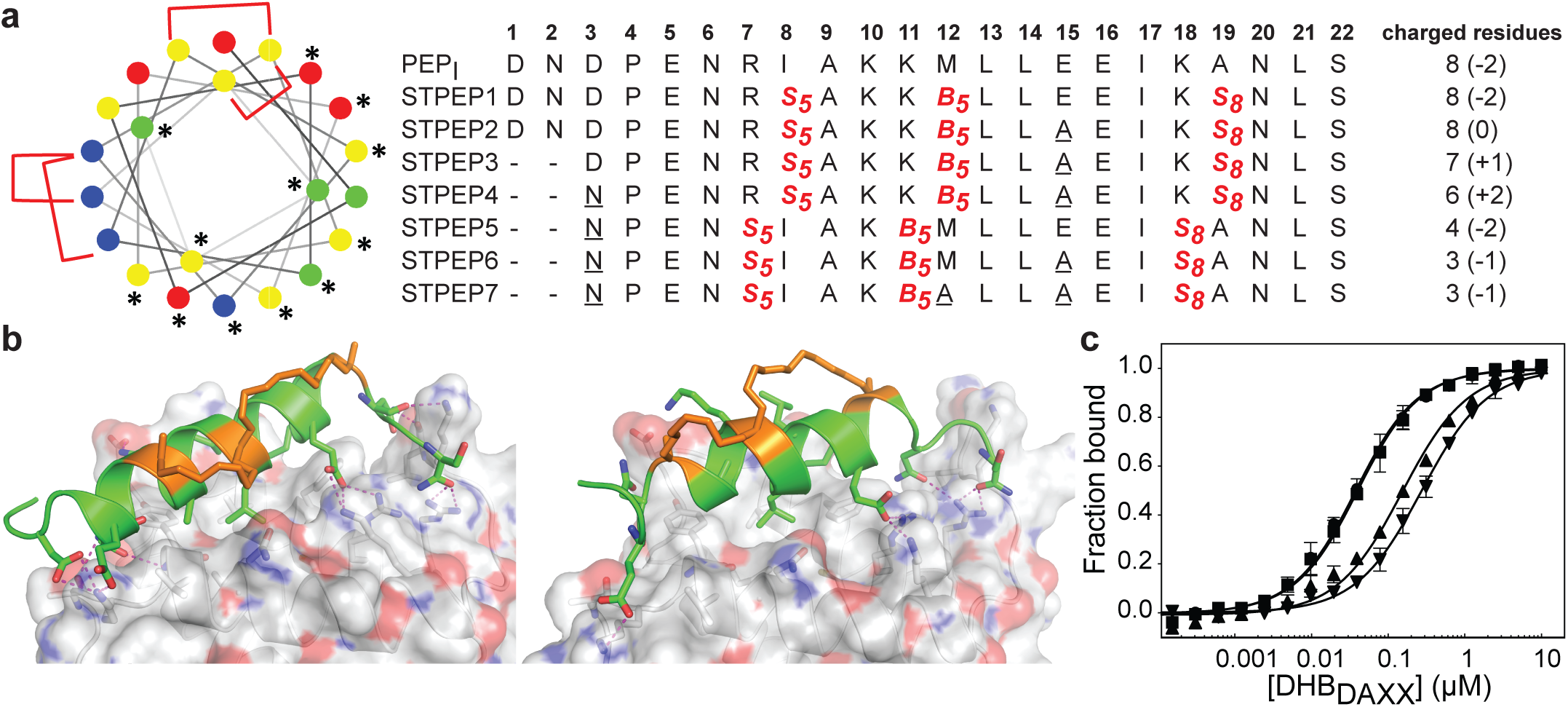
Design of stitched peptide inhibitors of DHB_DAXX_ PPI surface. A. Helical wheel representation of the linear PEP_I_ template sequence used for the design of stitched peptides. Annotations and colouring as in Fig 4 except that positions bridged by contiguous hydrocarbon staples i, i,i+4 plus i,i+7 are highlighted with red brackets. Sequences of the stitched peptide (STPEP1-7) designs are shown with stitch positions or mutated residues highlighted in red or underlined, respectively. Deleted residues are denoted ‘-’. Residue number is shown in bold above. The number of charged residues, indicating the hydrophobicity of each peptide, is shown to the right with the net charge in parentheses. **B** Structural models generated from MD snapshots for DHB_DAXX_ complexes with the two different stitch configurations. Structural representation as in Fig 5. Left: Stitched configuration (used in STPEP1-4) resulting from combining SPEP2 and SPEP7 staples. Right: Stitched configuration (used in STPEP5-7) resulting from combining SPEP1 and SPEP6 staples.

In HREMD simulations of STPEP1-7 peptides in solution starting from all alpha helical conformations, the stitched peptides were predicted to have comparable helicity to the parent singly stapled peptides. Also similar to the stapled peptides, per-residue helicity suggested that greater helicity is retained in the centre of the stitched peptides with increased flexibility at the termini (Figure S11). All stitched peptides remained stably bound to DHB_DAXX_ during the MD simulations of the STPEP/DHB_DAXX_ complexes with rmsd values reaching ∼3 Å or ∼2 Å from the corresponding starting conformations of DHB_DAXX_ or STPEPs, respectively (Figure S12). In addition, the hydrocarbon staples remained exposed to solvent without engaging the DHB_DAXX_ surface. Structural snapshots from the simulations of the complexes predicted that STPEP1, with a stitch based on SPEP2+7, retains all the interactions found in the individual stapled peptides, including those at the N- and C- termini, binding with similar affinity (Fig. 7B and Table S2). Sequential introduction of mutations E15A (STPEP2) and ΔD1−N2 (STPEP3) were also not found to affect binding. In a further attempt to reduce negative charge, a D3N mutant (STPEP4) was modelled, but this had a deleterious effect due to loss of a side-chain interaction between D3 and K122 of DHB_DAXX_ (Fig 7, Table S2). MD simulations of the peptides with a stitch based on SPEP 1+6 staples (STPEP5-7) predicted a further loss in binding affinity. Interestingly, reduced helicity in both bound and unbound forms of STPEP5-7 was observed, particularly towards the C-terminal region of the peptides, where a number of peptide-DHB_DAXX_ interactions were lost. This would not be expected from the i, i+7 staple incorporated into these peptides, which stabilises helix formation in this region in the singly stapled SPEP6 peptide (Fig. 5). Together, this suggests that the stitched peptides STPEP5-7 may not behave in the way expected from the individual staples.

We reasoned that a compromise between cell permeability and binding affinity of the stitched peptides might be required to target endogenous DAXX. Accordingly, we selected 4 stitched peptides with a range of properties to be synthesised based on our computer simulations; STPEP2, STPEP3, STPEP4 and STPEP7 (Fig 7A). These peptides were predicted to bind to DHB_DAXX_ with decreasing affinity as follows: STPEP2 ≅ STPEP3 > STPEP4 > STPEP7 (Table S2). In contrast, hydrophobicity of these peptides follows the reverse order, with STPEP7 being the most hydrophobic. We noted that the non-binding M12A mutation introduced in STPEP7 for ease of synthesis ironically slightly increased affinity in our simulations. It is not clear whether this resulted from the mutation or the staple position.

The selected stitched peptides were synthesised with an N-terminal FAM moiety to facilitate detection of binding to DHB_DAXX_, cell permeability and localisation. A marked decrease in solubilities of the stitched peptides was observed compared to the stapled peptides with limits of 11, 4, 2, or 49 μM determined for STPEP2, STPEP3, STPEP4 or STPEP7, respectively. This reduced solubility precluded the use of either ITC to determine the binding thermodynamics or CD to quantify helicity, but dissociation constants could be determined using fluorescence anisotropy experiments.

The trends in the dissociation constants obtained were in excellent agreement with the MD simulations confirming the validity of the computational analyses. No change in affinity was measured for either FAM-STPEP2 (E15A, Kd 32 nM) or STPEP3 (ΔD1-N2/E15A, Kd 35 nM) compared to FAM-SPEP7 (Kd 43 nM), whilst a ∼4 fold decrease in binding was measured for STPEP4 (ΔD1-N2/D3N/E15A, Kd 148 nM) and a ∼7 fold decrease was measured for STPEP7 (ΔD1-N2/D3N/E15A, Kd 269 nM), which contains the destabilising SPEP1+6 stitch (Figure 7D and Table S2).

### Cell permeability of stitched peptides

To test cellular permeability/localisation of the FAM-labelled stitched peptides, U2OS cells were chosen initially because they are negative for ATRX and are characterised by very large DAXX-associated nuclear PML bodies. It was therefore hoped that these could serve as a marker for targeting of the peptide/s to endogenous DAXX. Upon incubation of cells with FAM-labeled stitched peptides, a degree of endosomal escape into the cell body was observed, particularly with FAM-STPEP2 and FAM-STPEP7, (Fig. 8A) indicating an improvement in cytosolic localization of the peptide. Although no punctate nuclear foci were observed, with no measure of the absolute concentration/s within the cell it was not possible to estimate whether the amount of intracellular peptide was sufficient to bind to DAXX. To further estimate the extent of peptide intracellular delivery we used the NanoBRET assay.^51^ Chloroalkane versions of STPEP2 and STPEP7 were titrated in HT116 cells harbouring a cytosolic nanoluc-halotag sensor.^51^ While both peptides showed no significant cellular toxicity, Nanobret IC50s determined after 24 hrs incubation were 7.1 and 93 nM for STPEP2 and STPEP7, respectively (Fig 8B). Although not measured directly, we assumed the nuclear concentrations of peptide to be equal to that in the cytoplasm as it is well known that proteins smaller than 40 kDa can readily enter the nucleus by passive diffusion through nuclear pore complexes.^52^ Whilst these results are encouraging, the intracellular concentrations of the stitched peptides do not reach the concentrations suggested to be required by the binding affinities measured in vitro for STPEP2 and STPEP7 (32 nM and 269 nm respectively (Figure 7D and Table S2)). In conclusion, whilst the stitched peptides represent an improvement on the stapled peptides and display an increased cytosolic localization, the concentrations of peptides entering the cells appears to be too low to target endogenous DAXX and exert cellular activity. Taken together these results suggest that although our peptides were able to reach the cytosolic environment in absence of permanent membrane perturbation, future work will be required to improve cell permeability and establish whether the concentration of peptide within the cell/nucleus is sufficient to produce target engagement.

**Figure 8.**
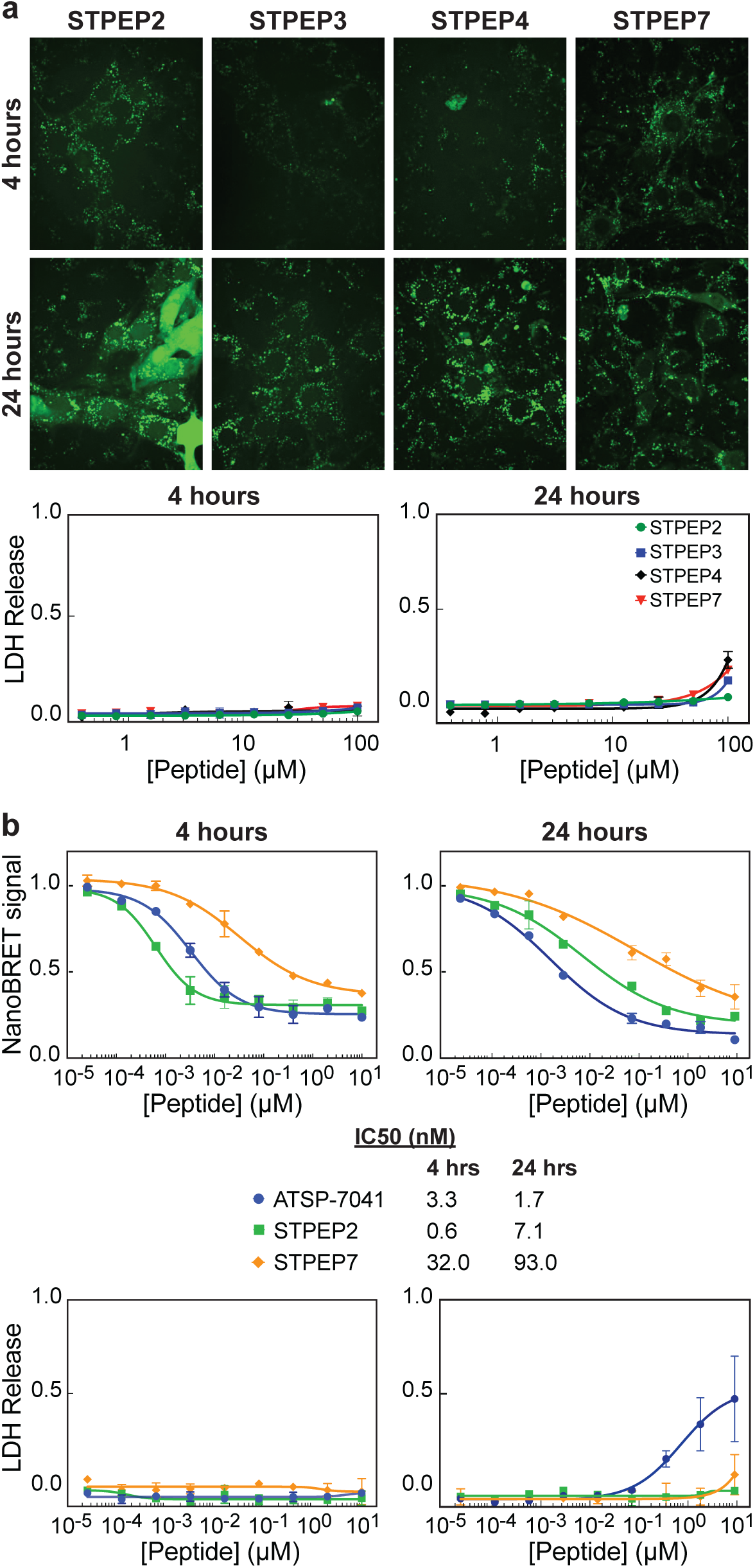
Cell permeability of stitched peptides. A. Micrographs: Representative confocal images of U2OS cells treated with 25 μM of the indicated FAM-labelled stitched peptides for either 4 or 24 hrs. LDH assays: Titrations of stapled peptides were performed on U2OS cells in presence of 2 % serum and LDH release was assessed at 4 and 24 hrs. B NanoBret assay: Titration of selected peptides vs Bret signal measured in HCT116 cells stably expressing the Nanoluc-Halotag fusion. The ATSP-7041 peptide was included as a positive control. Measurements were made 4 and 24 hrs after treatment with peptide and IC50s obtained from fitting the data are shown below the plots alongside the figure legends. LDH assays: Titrations of stitched peptides were performed on HCT116 cells in presence of 2 % serum and LDH release was assessed at 4 and 24 hrs. All data points in plots correspond to the mean of three independent measurements with errors corresponding to ± 1 SD.

## Conclusions

DAXX, a multifunctional protein, is frequently highly expressed in human cancers where it can promote cell proliferation (tumourigenesis) and chemoresistance.^53, 54^ In such cases DAXX is of considerable interest as a therapeutic target. With more than 70 reported protein interaction partners, elucidating the complex role/s of DAXX in cancer biology has proved challenging.

In this work, we hypothesized that targeting a promiscuous protein-protein interaction (PPI) surface of DAXX with peptidomimetic inhibitors could be a useful tool to decipher details of DAXX regulation and function and provide a starting point for exploiting DAXX as a therapeutic target. This PPI surface, located in the DAXX Helical Bundle domain (DHB_DAXX_, residues 55-144) interacts with multiple interaction partners in a mutually exclusive manner, marking it out as an important regulatory domain.^33, 36^ In addition, it has been reported that the N-terminal SIM of DAXX is sequestered by interaction with the same interaction surface, presumably to prevent promiscuous, unregulated interactions with non-cognate SUMOylated proteins that might otherwise occur.^35^ In turn, release of NSIM_DAXX_ through competitive inhibition might be expected to provide a means whereby DAXX could be sequestered in situations where it is aberrantly upregulated in cancer. Thus we also considered that our inhibitors might serve as a starting point for potential therapeutic development.

We report, through systematic evaluation of stapled and then stitched peptides based initially on the DAXX binding motif of ATRX, that it is possible to generate high affinity binders of the target protein-protein interaction surface. These stapled peptides bind with nM affinity to the DHB_DAXX_ (Table 1 and Fig. S7), and competition binding experiments show that such peptides can be displaced by DAXX binding peptides derived from either ATRX or P53, demonstrating specificity. Using a DAXX construct that includes both the DHB and NSIM of DAXX (residues 1-144 of DAXX), we also show that the same stapled peptide inhibitor can efficiently relieve self-inhibition, releasing NSIM_DAXX_ for binding to SUMO1 (Fig. 6 and Fig. S9).

Expecting that the release of NSIM_DAXX_ could have consequences for localisation of DAXX within the cell, we conducted cell-permeability experiments. After a degree of optimisation (addition of a second hydrocarbon linker and removal of non-essential charged residues) we obtained peptides that were cell permeable. However, the measured cytosolic concentrations were below the dissociation constants for peptide binding to DHB_DAXX_. Thus, further development of these peptide inhibitors, outside the scope of the results of the studies reported here, would likely be required to achieve the aim of targeting endogenous DAXX. This might include incorporating various different staples, mutations that vary the charge-hydrophobicity ratios of the peptides and/or conjugation with cyclic cell-penetrating peptides^55^ that will no doubt result in much more potent and cell active peptide inhibitors of DAXX. Nevertheless, this work provides an important starting point for the development of peptidomimetic tools to probe interactions mediated by DHB_DAXX_ in the cell that could also serve as therapeutic precursors.

## Materials and Methods

### Protein expression and purification

NSIM-DHB_DAXX_ (residues 1-144 hDAXX), DHB_DAXX_ (residues 55-144 hDAXX) and DID_ATRX_ (residues 1190-1325 hATRX) coding sequences were cloned into a pNIC28-bsa4 plasmid vector with a N-terminal His_6-_tev purification tag. GST-DID_ATRX_ and related fragments, were cloned into a pNIC-GST plasmid vector with a N-terminal His_6_-tev-GST tag. The SUMO-1 (residues 1-97 hSUMO1) coding sequence was synthesised and cloned into a pET28a plasmid with N-terminal His_6_-tev purification tag (supplied by Genscript Ltd).

All proteins were expressed in BL21(DE3) Rosetta T1R cells, cultured in 2xTY media with the exception of ^15^N-labelled NSIM-DHB_DAXX_ that was expressed in cells grown in M9 minimal media supplemented with 0.5 g/L ^15^NH_4_Cl. All proteins were initially purified after cell lysis by sonication using Nickel NTA affinity chromatography. A standard lysis/Ni-NTA binding buffer (20 mM HEPES pH 7.5, 0.3 M NaCl, 10 mM Imidazole, 1 mM DTT, 10 % glycerol) was used to load cell lysates and wash off unbound material, followed by a single step elution of bound proteins using the same buffer containing 0.3 M Imidazole. The N-terminal His-tag of the NSIM-DHB_DAXX_, DHB_DAXX_ or SUMO-1 proteins was cleaved off with tev protease during overnight dialysis of eluted proteins against Ni-NTA binding buffer at 4 °C. The cleaved tag, uncleaved protein, Tev protease and other contaminants were removed with Nickel NTA resin and the unbound protein purified further using a HiLoad 16/60 superdex 75 size exclusion column (GE Healthcare). His-GST-DID_ATRX_, GST- DID_ATRX_ fragments and His-DID_ATRX_ proteins were purified directly after Ni-NTA using HiLoad 16/60 superdex 200 or 75 size exclusion columns (GE Healthcare). With the exception of proteins used in NMR experiments, a standard storage buffer (20 mM HEPES pH 7.5, 0.3 M NaCl, 10 mM Imidazole, 1 mM DTT, 10 % glycerol) was used for the final size-exclusion and concentration steps prior to flash freezing of purified proteins. For ^15^N NSIM-DHB_DAXX_ and SUMO-1 proteins, phosphate buffer (20 mM sodium phosphate pH 6.5, 0.5 M KCl, 0.1 mM EDTA, 10 mM DTT) was used in the final purification steps. Protein concentrations were determined using the relevant molar extinction co-efficient at 280 nm.

### Peptide synthesis and purification

All non-fluorescent linear and stapled peptides (SPEP1- 7) were purchased from Mimotopes Pty Ltd. The FAM-SPEP7 peptide was purchased from Synpeptide Co Ltd. FAM- and chloroalkane labelled stitched peptides were synthesized in-house at the Institute of Chemical & Engineering Science, (A*STAR). Peptide purity was verified by HPLC and mass spectrometry. Further details of peptide purification can be found in supplementary methods. Unless otherwise described, concentrations of peptides lacking a FAM moiety or aromatic residue/s were determined using dilution factors from 10 mM peptide stocks in DMSO, where peptide stocks were prepared in accord with the synthesis report. Concentrations of FAM-labelled peptides were determined by measuring the absorbance at 495 nm, with extinction co-efficient of 75,000 cm^-1^ M^-1^. Concentrations of P53, Rassf1c and mdm2 peptides were determined using the relevant molar extinction co- efficient at 280 nm. All sequences and chemical modifications of the peptides used are detailed within main figures and/or supplementary data.

### Co-immunoprecipitation

Purified GST-tagged DID_ATRX_ and DID_ATRX_ fragments were immobilized on Glutathione sepharose 4B beads (GE Healthcare) under saturating conditions in binding buffer (20 mM HEPES, pH 7.5, 150 mM NaCl, 0.5 mM EDTA, 1 mM DTT, 10% Gycerol, 1% Triton). After removal and washing off (x3) excess GST-tagged proteins, Hela nuclear extract (Ipracell Ltd) in binding buffer supplemented with protease inhibitors (Roche Ltd) was added to the immobilized proteins. The samples were incubated at 4 °C for 4 hrs after which the beads were washed (x 4) with binding buffer. Bound proteins were eluted with 2 x SDS PAGE sample loading buffer (Thermo Fisher) and heating (95 °C, 5 min). After centrifugation, the samples was analysed by Western blot to detect full-length DAXX protein that had been immunoprecipitated. For Western blot analysis the immunoprecipitated proteins were separated by SDS PAGE (NuPAGE 4-12% Bis-Tris, Thermofisher), transferred to PVDF membrane, blocked in 10% BSA dissolved in PBS buffer containing 0.02 % Tween 20, probed with primary antibody (anti-DAXX, Sigma cat no. D7810, 1 in 8000 dilution) and detected with horseradish peroxidase-conjugated anti-rabbit secondary antibodies (CST, cat no 70745).

### Isothermal Titration Calorimetry (ITC)

ITC experiments were performed using a MicroCal PEAQ-ITC calorimeter at 25 °C (Malvern). All peptides and proteins were dialysed overnight against ITC binding buffer (20 mM HEPES, pH 7.5, 150 mM NaCl, 10 % Glycerol, 1mM DTT). The heat of binding was measured by 19 sequential 2 ml injections of peptide (or DID_ATRX_) at 0.1-1 mM into 0.2 ml DHB_DAXX_ at 0.01-0.1 mM. Data analysis and curve fitting to a one-site binding model was performed using the PEAQ-ITC analysis software package provided with the calorimeter. In the absence of an absorbance signal to measure the concentrations of dialysed peptides, the concentration of DHB_DAXX_ measured using A_280nm_ and the stoichiometry (n=1) were fixed during data fitting.

### Circular Dichroism Spectropolarimetry (CD)

Far UV CD spectra between 190 and 260 nm were measured with a 2 nm bandwidth in a 2 mm cuvette at 25 °C using a Chiroscan spectropolarimeter (Applied PhotoPhysics). Peptides at 0.05 mM were dialysed against 10 mM sodium phosphate, pH 7.5, 150 mM NaF at room temperature overnight. The raw ellipticity measurements were converted to Mean Residue Ellipticity (θ_MRE_) for plotting (Figure 4D and S3) and values of θ_MRE_ at 222 nm were used to calculate percentage helicity (Figure S4 and Table 2) using established methods.

### Fluorescence Polarisation

Fluorescence polarisation (FP) of the FAM-SPEP7 peptide with excitation and emission wavelengths of 485 nm and 535 nm respectively, bandwidths of 20 nm was measured using a Spark^TM^ 10M instrument (Tecan) at 25 °C. A constant concentration of 10 nM FAM-SPEP7 was maintained throughout both saturation and competition binding experiments. In saturation binding experiments characterised by increasing FP upon binding, FAM-SPEP7 was titrated with varying amounts of either DHB_DAXX_ or NSIM-DHB_DAXX_ proteins. For competition binding experiments, a constant concentration of DHB_DAXX_ (0.75 mM) was maintained to generate an 80% saturated FAM- SPEP7/DHB_DAXX_ complex in the absence of competitor. Displacement of the FAM-SPEP7 peptide from this complex, characterised by decreasing FP was quantified by titration with increasing amounts of competitor ligand. All titrations were performed by making a 1:1 serial dilution of titrant into solutions of FAM-SPEP7 or FAM-SPEP7/DHB_DAXX_ complex and allowing samples to equilibrate for 30 min at room temperature prior to measurement. FP data were fitted to a two-state binding model for saturation binding experiments or a competitive binding model for competition binding experiments.^56^

### 1H-15N HSQC NMR Spectroscopy

^1^H-^15^N HSQC NMR spectra were recorded using a cryo-probe equipped Bruker Avance II 600 MHz spectrometer, processed and analysed using TopSpin4.0 software. Proteins were transferred into NMR buffer (10 mM potassium phosphate, pH 6.5, 100 mM KCl, 0.1 mM EDTA, 10 mM DTT, 10% D_2_O) and concentrated prior to titration. We first titrated ^15^N DHB_DAXX_ (100 mM in NMR buffer) with FAM-SPEP7 (10 mM in DMSO) to make a saturated complex. We observed discrete changes in the ^1^H-^15^N HSQC spectrum during the titration, where the transition between unbound and bound forms was in slow exchange on the NMR timescale (Figure S9). A small quantity of precipitated complex was formed during the titration. This was removed and the final concentration of soluble ^15^N DHB_DAXX_/FAM-SPEP7 complex was determined using the absorbance at 280 nm. This complex was titrated with unlabelled SUMO-1 (1.7 mM in NMR buffer), upon which we saw discrete changes in the ^1^H-^15^N HSQC spectrum, where affected cross peaks were in intermediate or fast exchange. We followed 9 fast exchange peaks during the titration to determine the saturation point of the observed transition. The chemical shift changes observed for each peak were plotted as a function of the ligand-substrate ratio and fitted to a 1:1 binding model to determine the individual dissociation constants (Figure S9), using the following equation:

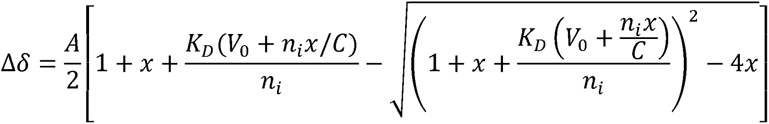

where, *A* = Normalization constant, *x* = ligand-substrate molar ratio, *K_D_* = dissociation constant, *V_0_*= initial volume (in µl), *n_i_* = initial amount of DHB_DAXX_—FAM-SPEP7 complex (in nmol), *C* = stock concentration of ligand (SUMO-1, in mM).

### LDH assays Lactate (LD) Dehydrogenase Release Assay

Cells were seeded into a 96-well plate in DMEM containing 10% FCS at a cell density of 5000 cells per well and incubated for 24 hours prior to replacement of cell media with 90 µl of culturing medium with 2% FCS.

Cells were then treated with 10 ul of peptide for either 4 or 24 hrs. Final working concentration of DMSO was 1% v/v. Corresponding negative control wells with 1% DMSO only were also prepared. Cytosolic lactate dehydrogenase release was detected using the cytoTox 96® non-radioactive cytotoxicity Assay kit (Promega) as per manufacturer’s instructions. Measurements were carried out using an Envision multiplate reader (Perkin- Elmer). Maximum LDH release was defined as the amount of LDH released when cells were lysed in the presence of 0.1% TRITON X-100 and was used to normalize the results.

### Live Cell Confocal Microscopy

100,000 U20S cells were plated in 3.5 cm glass bottomed dishes in DMEM containing 10% FCS. After 24h, the media was replaced with DMEM containing 0% or 2% FBS and treated with 12.5 μM of indicated FAM-labelled peptides for 5 hrs or 24 hrs. Cells were then washed twice with PBS saline and media replaced with Optimem without phenol red. Confocal images were acquired at the indicated time points using a Yokogawa CSU-22spinning disk confocal built around a motorized Nikon Eclipse Ti microscope equipped with a stage-top incubator and CO2control system, equipped with a 100 × 1.4 NA Plan Apo objective lens, photometrics Cool SNAPHQ2 camera and 491nm laser. Acquisition parameters, shutters, filter positions and focus were controlled by MetaMorph software(Molecular Devices)).

### NanoBRET assay

HCT116 stably expressing Nanoluc-Halotag fusion were seeded for overnight incubation at 60,000 cells/well in a 96-well white opaque tissue culture plate. The day after medium was replaced with 90 ul of Optimem without red phenol in presence or absence of 2% FCS. Cells were then titrated with chloroalkane-labelled stitched peptides and incubated for either 4 or 24 hrs at 37 °C, 5% CO2. Final working concentration of DMSO was 1% v/v. Labelling of unoccupied Nanoluc-Halotag molecules was performed 30 min before each end point by adding 10 ul of 618 Nanobret-Ligand solution 12x per well. To measure BRET signal inhibition, 10 ul of 12x Nano-Glo Substrate solution was added to each well and plates were read on the Envision instrument immediately.

### Computational Modelling

A model of the three dimensional atomistic structure of DID_ATRX_ (residues from 1190-1325) was constructed using the program I-TASSER^40^ and the atomistic structure of the complex between the experimental structure (see below) of DHB_DAXX_ and the structure of PEP_ATRX_ extracted from the DID_ATRX_ model (above) was constructed using the program HADDOCK (High Ambiguity Driven biomolecular DOCKing).^41, 42^ The available experimental structures of DHB_DAXX_ in apo form (PDB 2KZS), in complex with Rasf1c (PDB 2KZU) and in complex with PEP_ATRX_ (PDB 5GRQ, resolution 1.5 Å) were used to design stapled and stitched peptide inhibitors of DAXX protein. The Xleap module of the Amber18 program ^57^ was used to prepare the system for the Molecular Dynamics (MD) simulations. The parameters for the staple and stitched linkers were taken from our previous study.^58^ All simulation systems were neutralized with an appropriate number of counter ions. The neutralized system was solvated in an octahedral box with TIP3P^59^ water molecules, leaving at least 10 Ǻ between the solute atoms and the borders of the box. MD simulations were carried out with the pmemd module of the Amber18 package, using the ff14SB force field.^60^ All MD simulations were carried out in explicit solvent at 300 K unless specified. During all the simulations, the long-range electrostatic interactions were treated with the particle mesh Ewald ^61^ method using a real space cut off distance of 9 Ǻ. The settle^62^ algorithm was used to constrain bond vibrations involving hydrogen atoms, which allowed a time step of 2 fs to be used during the simulations. Solvent molecules and counter ions were initially relaxed using energy minimization (1000 steps of Steepest Descent minimizer) with restraints (force constant: 50 kcal mol^-1^ Å^-2^) on the protein and peptide atoms. This was followed by unrestrained energy minimization to remove any steric clashes. Subsequently the system was gradually heated from 0 to 300 K using MD simulations with positional restraints (force constant: 50 kcal mol^-1^ Å^-2^) on protein and peptides over a period of 0.25 ns allowing water molecules and ions to move freely followed by gradual removal of the positional restraints and a 2 ns unrestrained equilibration at 300 K. The resulting systems were used as starting structures for the respective production phase of the MD simulations. For each case, three independent (using different initial random velocities) MD simulations were carried out starting from the well equilibrated structures. Each MD simulation was carried out for 100 ns and conformations were recorded every 4 ps. To enhance the conformational sampling, each of these peptides were subjected to Biasing Potential Replica Exchange MD (BP-REMD) simulations. The BP-REMD technique is a type of Hamiltonian REMD method where a biasing potential that promotes dihedral transitions along the replicas is used.^63, 64^ For each system, BP-REMD was carried out with eight replicas including a reference replica without any bias. BP-REMD was carried out for 50 ns with exchange between the neighbouring replicas attempted for every 2 ps and accepted or rejected according to the metropolis criteria. Conformations sampled at the reference replica (no bias) was used for further analysis. Simulation trajectories were visualized using VMD^65^ and figures were generated using Pymol.^66^

### Binding Energy calculations and energy decomposition analysis

The binding energies between the peptides and their partner proteins were calculated using the Molecular Mechanics Poisson Boltzmann Surface Area (MMPBSA) method.^67, 68^ A set of 250 conformations, extracted every 200ps, from the last 50 ns of the simulations were used for the binding energy calculations. Entropy calculations are computationally intensive and do not converge easily and hence are ignored. The effective binding energies were decomposed into contributions of individual residues using the Generalized Born approximation to the MMPBSA method (called MMGBSA) energy decomposition scheme. The MMGBSA calculations were carried out in the same way as in the MMPBSA calculations. The polar contribution to the solvation free energy was determined by applying the generalized born (GB) method (igb =2)^57^, using mbondi2 radii. The non-polar contributions were estimated using the ICOSA method^57^ by a solvent accessible surface area (SASA) dependent term using a surface tension proportionality constant of 0.0072 kcal/mol Å^2^. The contribution of peptide residues was additionally explored by carrying out in-silico alanine scanning in which each peptide residue is mutated to alanine in each conformation of the MD simulation and the change in the binding energy relative to that of the wild-type peptide is calculated using MMPBSA and averaged over all the conformations.

## Acknowledgements

We thank staff at the NTU Institute of Structural Biology (NISB), Nanyang Technological University, for providing access to facilities; The Singapore Eye Research Institute for access to facilities; Charles Johannes and staff at the Institute of Chemical & Engineering Science (A*STAR) for synthesis of stitched-peptides; The Bio-informatics Institute A*STAR and The National Supercomputing Centre (NSCC) for computational support. This work has been supported by NTU Lee Kong Chian School of Medicine and the Singapore Ministry of Education (MOE) Academic Research Fund (AcRF) Tier 3 (MOE2012-T3-1-001) grant.

## Author contributions

C.J., S.K and C.S.V. conceptualized and designed the study. C.J purified all protein reagents and performed all in vitro binding assays, with assistance from F.W for the collection of NMR data. C.J. and F.W processed and analysed NMR spectra. Y.F performed live-cell imaging, LDH and NanoBret assays with assistance from SRR. Y.F performed anaylsis of all cell-based assays. S.K performed all computational modelling and analysis. R.L assisted with interpretation of CD data. C.J. wrote the paper with input from S.K, C.S.V and D.R. All authors contributed to the interpretation of the data as applicable to their technical contribution.

