## Supplementary methods and figure legends for "Stitched peptides as potential cell permeable inhibitors of oncogenic DAXX protein"

**Synthesis of FAM- and Halo-tagged stitched peptides**

*General information:* RAMAGE resin was obtained from PCAS Biomatrix. Fmoc-amino acids, Hexafluorophosphate Azabenzotriazole Tetramethyl Uronium (HATU), and 1-Hydroxy-7-azabenzotriazole (HOAt) were obtained from Advanced Chemtech (Louisville, KY). Trifluoroacetic acid (TFA) and N, N-Diisopropylethylamine (DIPEA) was purchased from Tokyo Chemical Industry (Tokyo, Japan) while all other solvents and reagents were obtained from Fisher Scientific (Loughborough, United Kingdom). All reagents were used as received.

*For FAM-labelled peptides:* 5-carboxy fluorescein was purchased from Beijing Okeanos Tech (Beijing, China).

*For Halotagged peptides:* Halotag was purchased from Beijing Okeanos Tech (Beijing, China).

*Solid-phase peptide synthesis:* The linear peptide was synthesized on the CEM peptide synthesizer. 1 M OxymaPure with 0.1 M DIPEA in DMF was used as the coupling agent and 0.5 M DIC in DMF was the activating agent. Fmoc deprotection was done with 20% Piperidine in DMF. Following ring closing metathesis, 5-carboxy fluorescein or Halotag was coupled onto the N-terminus manually with pre-activated solutions (7 min) of 2.8 eq. HATU, 3 eq. HOAt, and 6 eq. DIPEA for 1 h.

*Ring closing metathesis:* 0.2 equiv. Grubbs I catalyst was dissolved in DCE (5 mg/mL) was bubbled in the resin for four times for two hours each.

*Cleavage:* Following synthesis, the peptide resin was shrunk and dried with washes of methanol and diethyl ether. It was cleaved from the resin with a cleavage cocktail of TFA/Tips/H2O (95:2.5:2.5) for 2 h. The resin was filtered off and peptide precipitated in diethyl ether (50 mL). The peptide was re-dissolved in a mixture of ACN/H2O (1:1) and lyophilised.

*Peptide purification and analysis:* The dried peptide was re-dissolved in acetonitrile and water (1:1) and purified *via* reverse-phase HPLC using an Agilent 1260 Infinity system fitted with a Phenomenex^®^ preparative column (Jupiter C12, 4 µm, Proteo 90 Å, 250 x 10 mm). Eluents used were 0.1% aqueous TFA in water and 0.1% TFA in acetonitrile. Peptide purity and molecular weight were confirmed *via* UPLC-MS using an Agilent 1260 Infinity II system fitted with a Phenomenex® analytical column (Aeris 1.7 µm, Peptide XB-C18 100 LC Column, 150 x 2.1 mm). Eluents used were 0.1% aqueous formic acid in water and 0.1% formic acid in acetonitrile.

**Supplementary figures:**

**Figure S1.** **ITC isotherms for experiments corresponding to thermodynamic binding parameters listed in Table 1.**

**Figure S2. Stability of docked DHB_DAXX_/PEP_RassF1C_ and DHB_DAXX_/PEP_ATRX_ complexes. A** Cartoon representations showing 7 poses of the PEP_ATRX_ helix (multiple colours) docked with DHB_DAXX_ (grey). **B** RMSD of conformations of PEP_Rassf1C_ sampled during MD simulations starting from the DHB_DAXX_/PEP_Rassf1C_ solution structure. **C** RMSD of conformations of PEP_ATRX_ peptides sampled during MD simulations starting from different docked poses of PEP_ATRX_; (left) RMSD of the peptides with stable binding; (right) RMSD of peptides with unstable binding during MD simulations.

**Figure S3. Structure and stability analysis of PEP_ATRX_ and DHB_DAXX_ in apo and bound forms. A** Time evolution of secondary structures in apo PEP_ATRX_ sampled during BP-REMD simulations (blue: α - helix, gray: 3_10_- helix, yellow: turn, green: bend, white: coil). **B** CD spectrum of PEP_ATRX_ showing % helicity. **C** Probability distributions of RMSD of conformations sampled during simulations; (left) apo DHB_DAXX_; (middle) bound DHB_DAXX_ from DHB_DAXX_/PEP_ATRX_ complex, and (right) bound PEP_ATRX_ from DHB_DAXX_/PEP_ATRX_ complex. Black and red corresponds to RMSD of conformations with or without the flexible N- and/or C- terminal residues described in the main text, respectively.

**Figure S4. Conformational analysis of stapled peptides.** Time evolution of the secondary structure of SPEP1 to 7 during BP-REMD simulations (blue: α - helix, grey: 3_10_- helix, yellow: turn, green: bend, white: coil, red: beta strand). The overall peptide conformations sampled were used to calculate percentage helicity (‘Calc’). CD spectra of peptides, fitted to obtain experimental estimates of percentage helicity (‘Exp’) are plotted alongside. Experimental data and fitted curves are coloured purple and green, respectively.

**Figure S5. Computational analysis of the stability of DHB_DAXX_/SPEP complexes.** RMSD of conformations sampled during MD simulations of DHB_DAXX_/SPEP complexes; (top) DHB_DAXX_ and (bottom) stapled peptides (SPEP). The different colors, black, red, green, blue, yellow, brown and grey corresponds to different DHB_DAXX_ /SPEP1 to 7 complexes in numerical order.

**Figure S6: Per-residue contributions to binding of DHB_DAXX_ by SPEP1-7 calculated from MD simulations.** Binding free energies of individual SPEP residues to DHB_DAXX_ were calculated using the MMPBSA approach (see Methods). The staple linker positions are highlighted in blue.

**Figure S7. ITC data for binding of DHB_DAXX_ to SPEP1 to 7.**

**Figure S8. Competitive displacement of FAM-SPEP7 from DHB_DAXX_ by non-fluorescent SPEP peptides 1-7.** Binding data with standard error bars derived from three independent measurements are shown for each peptide (closed symbols). Curves obtained from fitting the data to a competitive binding model (solid black line) are shown with simulated curves using Kd values obtained from ITC data (dashed red line) included for comparative purposes.

**Figure S9. Formation of the ^15^N-NSIM-DHB_DAXX_/FAM-SPEP7 pre-complex and subsequent binding of SUMO-1 in sequential titrations monitored using NMR. A.** Overlaid ^1^H-^15^N HSQC spectra of unbound ^15^N-NSIM-DHB_DAXX_ (blue) and a saturated complex formed with FAM-SPEP7 (red) are shown. The transition of a single resonance projected in 1D (inset) with an additional mid-titration point is included (black), indicating slow exchange. **B.** (Top) Three panels showing overlaid sections of the ^1^H-^15^N HSQC spectra of ^15^N-NSIM-DHB_DAXX_/FAM-SPEP7 in the absence (black) and presence (grey) of increasing quantities of SUMO-1. Multi-coloured arrows indicate 9 individual resonances whose discrete chemical shift changes, in fast exchange, were measured during the titration. (Bottom) Titration curves of the 9 resonances, coloured as indicated above. The reported Kd value is the mean ± SD of the 9 individually fitted curves. Combined amide chemical shift changes, Dd (ppm), were calculated as ((Dd_1H_)^2^+(0.2Dd_15N_)^2^)^1/2^.

**Figure S10. Cytotoxicity and cell permeability data SPEP1-7. A** HCT116 cells were titrated with peptides in the presence of 2 % serum and LDH release was assessed after 4 hrs incubation. **B** Cellular uptake was assessed by live cell imaging after treating HCT116 cells for 4 hrs with 25 uM of FAM-SPEP7 in the same conditions.

**Figure S11. Conformational analysis of stitched peptides.** Time evolution of the secondary structure of STPEP1 to 7 during BP-REMD simulations (blue: α - helix, grey: 3_10_- helix, yellow: turn, green: bend, white: coil, red: beta strand). The overall peptide conformations sampled were used to calculate percentage helicity for each peptide.

**Figure S12. Computational analysis of the stability of DHB_DAXX_/STPEP complexes.** RMSD of conformations sampled during MD simulations of DHB_DAXX_/STPEP complexes; (top) DHB_DAXX_ and (bottom) stapled peptides (SPEP). The different colors; black, red, green, blue, yellow, brown and grey, correspond to different DHB_DAXX_ /STPEP1 to 7 complexes in numerical order.

**Supplementary figures**

**Figure S1.** **ITC isotherms for experiments corresponding to thermodynamic binding parameters listed in Table 1.**

**Figure S2. Stability of docked DHB_DAXX_/PEP_RassF1C_ and DHB_DAXX_/PEP_ATRX_ complexes. A** Cartoon representations showing 7 poses of the PEP_ATRX_ helix (multiple colours) docked with DHB_DAXX_ (grey). **B** RMSD of conformations of PEP_Rassf1C_ sampled during MD simulations starting from the DHB_DAXX_/PEP_Rassf1C_ solution structure. **C** RMSD of conformations of PEP_ATRX_ peptides sampled during MD simulations starting from different docked poses of PEP_ATRX_; (left) RMSD of the peptides with stable binding; (right) RMSD of peptides with unstable binding during MD simulations.

**Figure S3. Structure and stability analysis of PEP_ATRX_ and DHB_DAXX_ in apo and bound forms. A** Time evolution of secondary structures in apo PEP_ATRX_ sampled during BP-REMD simulations (blue: α - helix, gray: 3_10_- helix, yellow: turn, green: bend, white: coil). **B** CD spectrum of PEP_ATRX_ showing % helicity. **C** Probability distributions of RMSD of conformations sampled during simulations; (left) apo DHB_DAXX_; (middle) bound DHB_DAXX_ from DHB_DAXX_/PEP_ATRX_ complex, and (right) bound PEP_ATRX_ from DHB_DAXX_/PEP_ATRX_ complex. Black and red corresponds to RMSD of conformations with or without the flexible N- and/or C- terminal residues described in the main text, respectively.

**Figure S4. Conformational analysis of stapled peptides.** Time evolution of the secondary structure of SPEP1 to 7 during BP-REMD simulations (blue: α - helix, grey: 3_10_- helix, yellow: turn, green: bend, white: coil, red: beta strand). The overall peptide conformations sampled were used to calculate percentage helicity (‘Calc’). CD spectra of peptides, fitted to obtain experimental estimates of percentage helicity (‘Exp’) are plotted alongside. Experimental data and fitted curves are coloured purple and green, respectively.

**Figure S5. Computational analysis of the stability of DHB_DAXX_/SPEP complexes.** RMSD of conformations sampled during MD simulations of DHB_DAXX_/SPEP complexes; (top) DHB_DAXX_ and (bottom) stapled peptides (SPEP). The different colors, black, red, green, blue, yellow, brown and grey corresponds to different DHB_DAXX_ /SPEP1 to 7 complexes in numerical order.

**Figure S6: Per-residue contributions to binding of DHB_DAXX_ by SPEP1-7 calculated from MD simulations.** Binding free energies of individual SPEP residues to DHB_DAXX_ were calculated using the MMPBSA approach (see Methods). The staple linker positions are highlighted in blue.

**Figure S7. ITC data for binding of DHB_DAXX_ to SPEP1 to 7.**

**Figure S8. Competitive displacement of FAM-SPEP7 from DHB_DAXX_ by non-fluorescent SPEP peptides 1-7.** Binding data with standard error bars derived from three independent measurements are shown for each peptide (closed symbols). Curves obtained from fitting the data to a competitive binding model (solid black line) are shown with simulated curves using Kd values obtained from ITC data (dashed red line) included for comparative purposes.

**Figure S9. Formation of the ^15^N-NSIM-DHB_DAXX_/FAM-SPEP7 pre-complex and subsequent binding of SUMO-1 in sequential titrations monitored using NMR. A.** Overlaid ^1^H-^15^N HSQC spectra of unbound ^15^N-NSIM-DHB_DAXX_ (blue) and a saturated complex formed with FAM-SPEP7 (red) are shown. The transition of a single resonance projected in 1D (inset) with an additional mid-titration point is included (black), indicating slow exchange. **B.** (Top) Three panels showing overlaid sections of the ^1^H-^15^N HSQC spectra of ^15^N-NSIM-DHB_DAXX_/FAM-SPEP7 in the absence (black) and presence (grey) of increasing quantities of SUMO-1. Multi-coloured arrows indicate 9 individual resonances whose discrete chemical shift changes, in fast exchange, were measured during the titration. (Bottom) Titration curves of the 9 resonances, coloured as indicated above. The reported Kd value is the mean ± SD of the 9 individually fitted curves. Combined amide chemical shift changes, Dd (ppm), were calculated as ((Dd_1H_)^2^+(0.2Dd_15N_)^2^)^1/2^.

**Figure S10. Cytotoxicity and cell permeability data SPEP1-7. A** HCT116 cells were titrated with peptides in the presence of 2 % serum and LDH release was assessed after 4 hrs incubation. **B** Cellular uptake was assessed by live cell imaging after treating HCT116 cells for 4 hrs with 25 uM of FAM-SPEP7 in the same conditions.

**Figure S11. Conformational analysis of stitched peptides.** Time evolution of the secondary structure of STPEP1 to 7 during BP-REMD simulations (blue: α - helix, grey: 3_10_- helix, yellow: turn, green: bend, white: coil, red: beta strand). The overall peptide conformations sampled were used to calculate percentage helicity for each peptide.

**Figure S12. Computational analysis of the stability of DHB_DAXX_/STPEP complexes.** RMSD of conformations sampled during MD simulations of DHB_DAXX_/STPEP complexes; (top) DHB_DAXX_ and (bottom) stapled peptides (SPEP). The different colors; black, red, green, blue, yellow, brown and grey, correspond to different DHB_DAXX_ /STPEP1 to 7 complexes in numerical order.
