## Supplementary figures and tables for "Stitched peptides as potential cell permeable inhibitors of oncogenic DAXX protein"

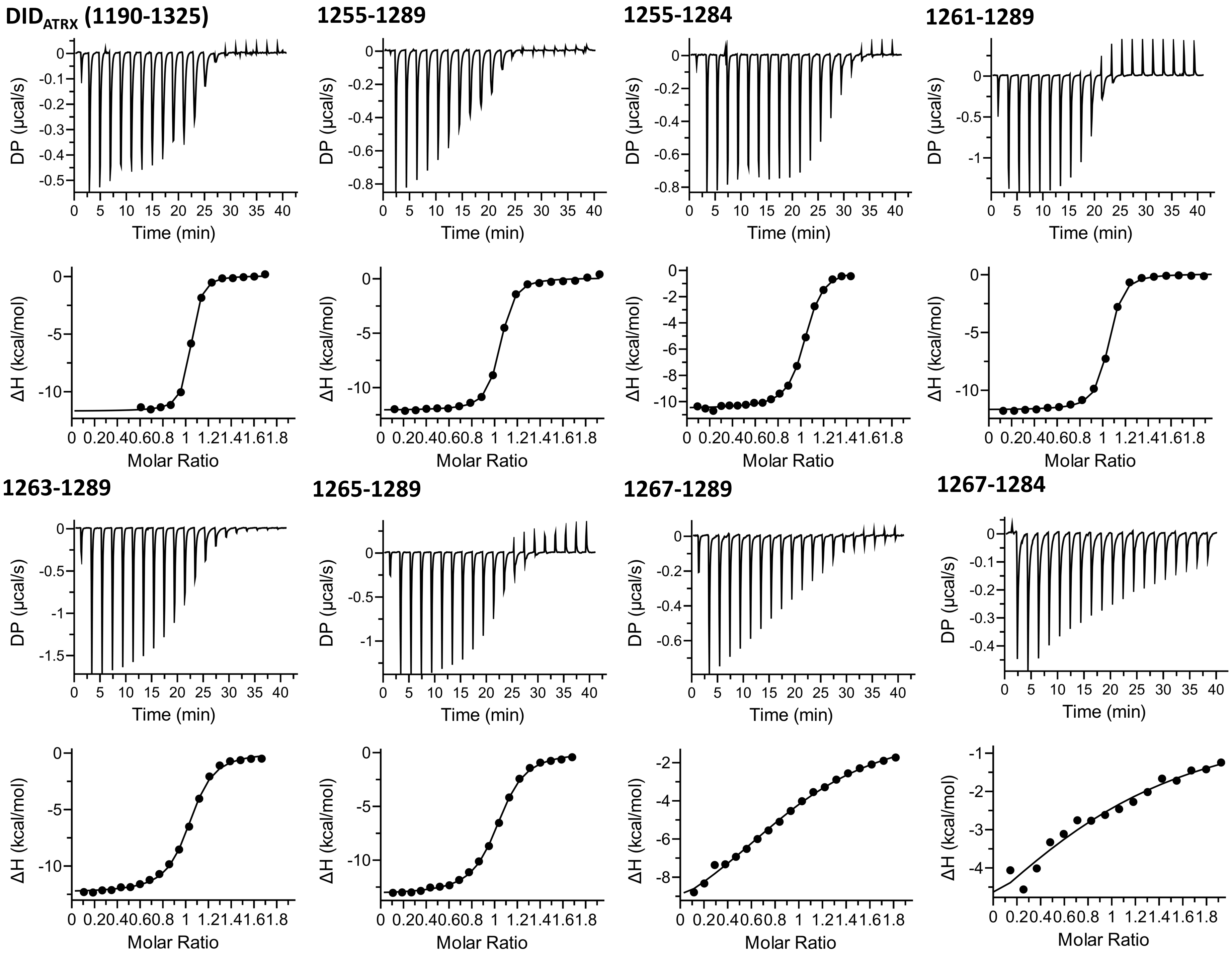


**Figure S1. ITC isotherms for experiments corresponding to thermodynamic binding parameters listed in Table 1.**

**Figure S2. Stability of the docked $DHB_{DAXX}/PEP_{RassF1C}$ and $DHB_{DAXX}/PEP_{I}$ complex.**

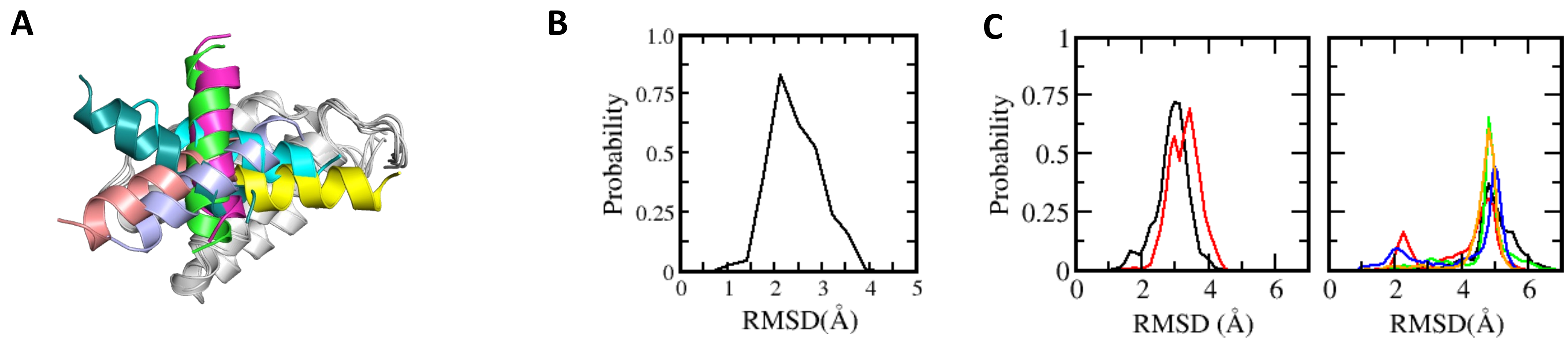

**Figure S3. Structure and stability analysis of $PEP_I$ and $DHB_{DAXX}$ in apo and bound forms.**

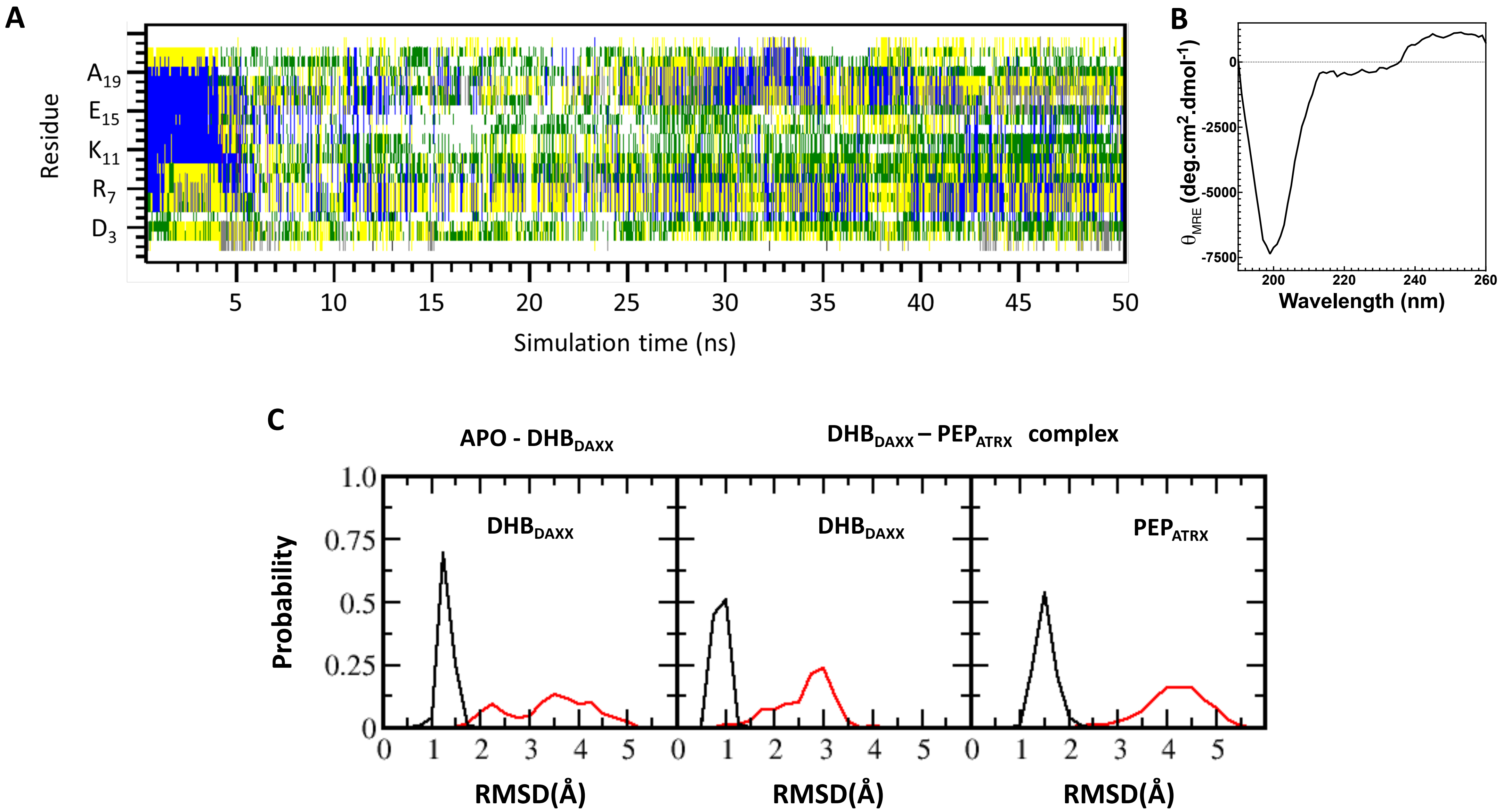

### Figure S4. Conformational analysis of stapled peptides.

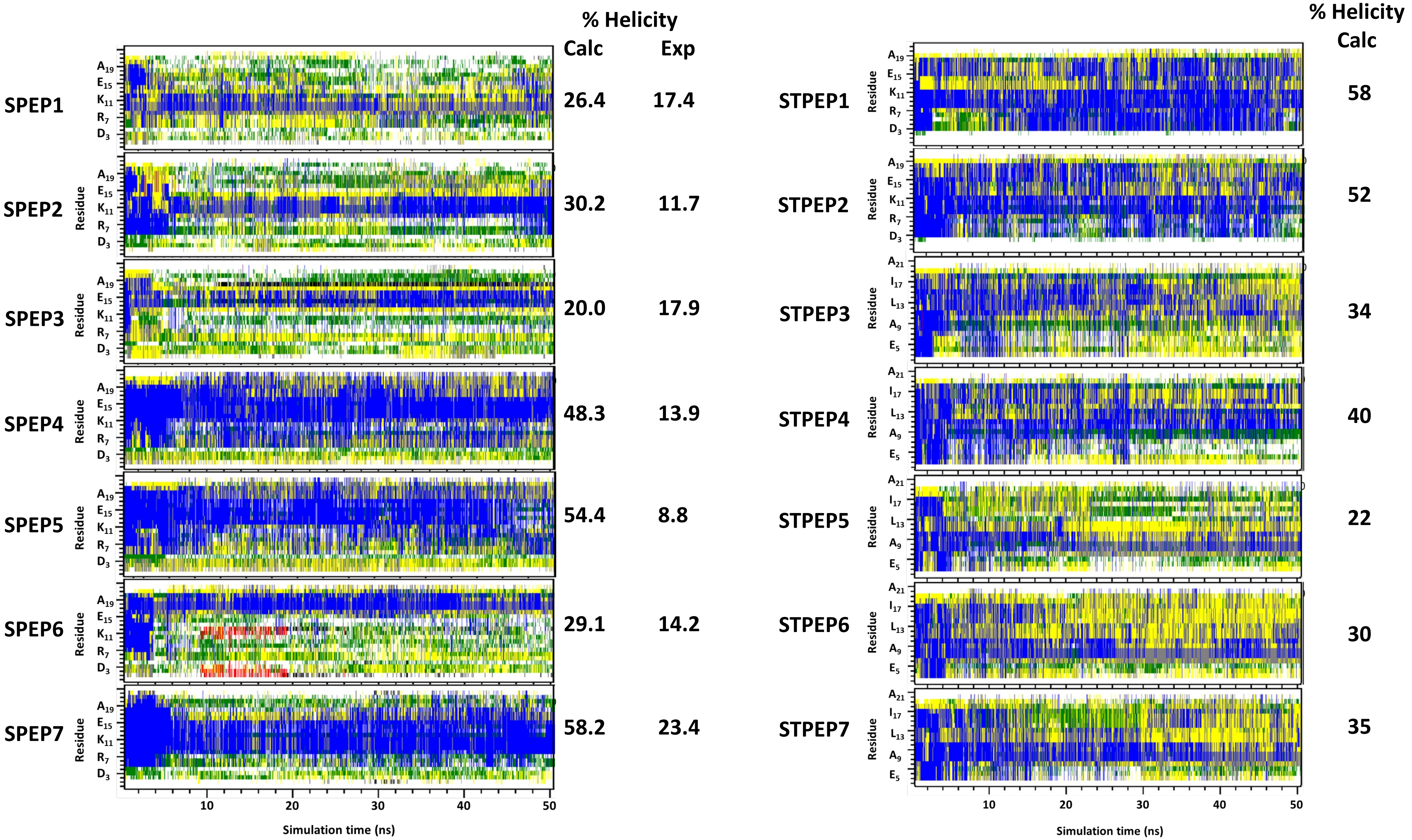

### Fig S5. Computational analysis of the stability of $DHB_{DAXX}$/SPEP complexes.

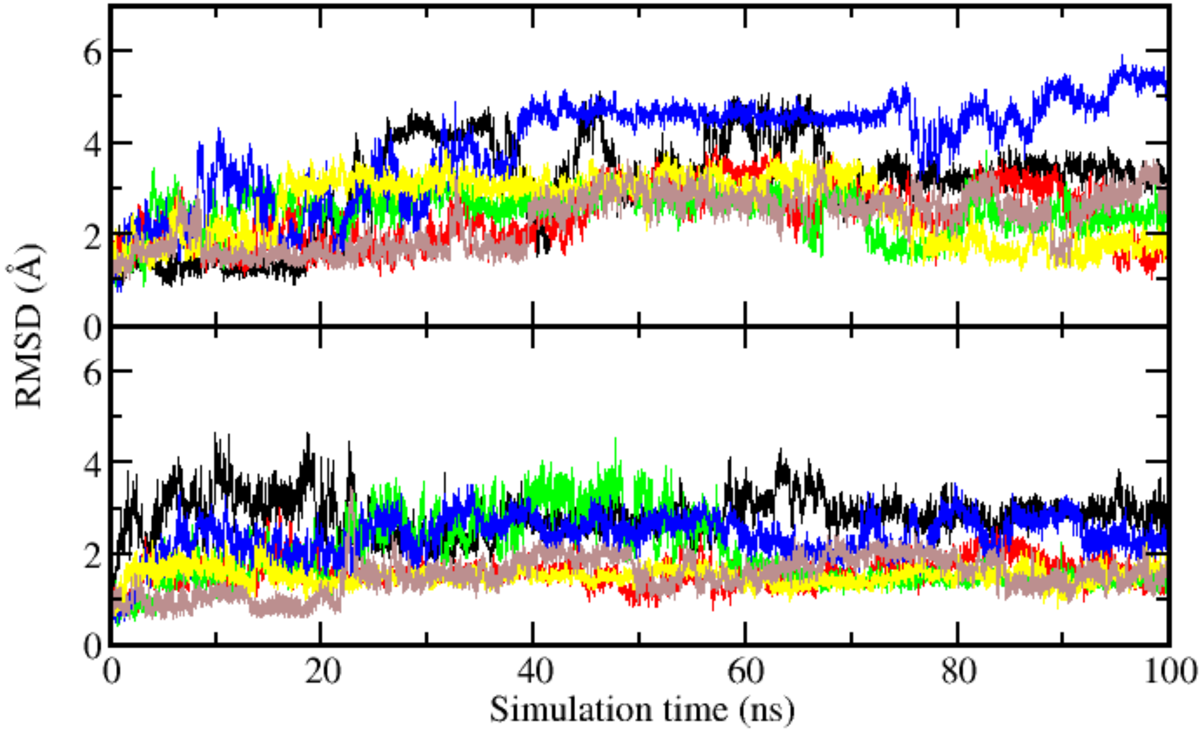

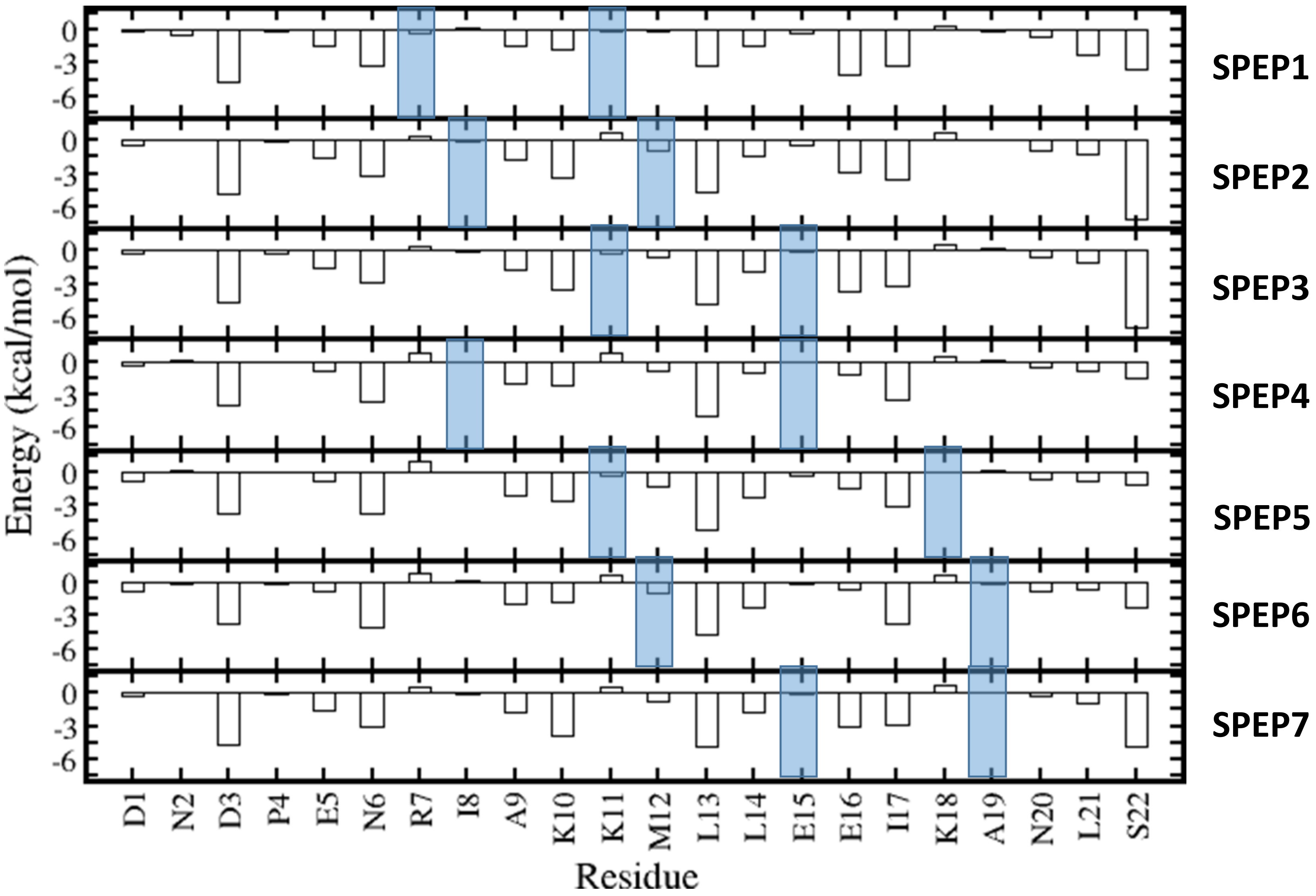


**Figure S6: Per-residue contributions to binding of $DHB_{DAXX}$ by SPEP1-7 calculated from MD simulations.**

**Table S1. Binding parameters for the formations of $DHB_{DAXX}$/SPEP complexes determined through computational methods$^{\$}$**

| Peptide | Ele | Vdw | Polar | Nonpolar | ΔG* |
|---|---|---|---|---|---|
| $PEP_I$ | -836.9 (40) | -66.2 (1.4) | 829.2 (33.1) | -12.7 (0.2) | -84.1 (7.1) |
| SPEP1 | -929.4 (41) | -73.1 (1.4) | 931.5 (42) | -13.5 (0.2) | -84.3 (2.3) |
| SPEP2 | -850.5 (13) | -67.1 (0.6) | 834.8 (8) | -12.5 (0.3) | -92.1 (1.8) |
| SPEP3 | -853 (1.1) | -67.9 (2.3) | 839.8 (7) | -12.7 (0.7) | -82.4 (3.3) |
| SPEP4 | -564.4 ( 66) | -69.3 (1.6) | 877 (31) | -11.9 (0.5) | -73.7 (3.3) |
| SPEP5 | -892.6 (26) | -71.7 (7.2) | 649 (67) | -11.3 (0.8) | -70.9 (5.5) |
| SPEP6 | -641.1 (74) | -70.3 (3.6) | 721 (25) | -12.3 (0.1) | -80.1 (4.0) |
| SPEP7 | -726 (31) | -68.1 (1.1) | 574 (64) | -11.2 (0.3) | -86.3 (1.4) |

**$^{\$}$All values are calculated in kcal mol$^{-1}$ at 300K. Errors of 1 SD are shown in parentheses. *Entropy changes are not calculated, therefore the binding energy calculated here corresponds to only the enthalpy contribution.**

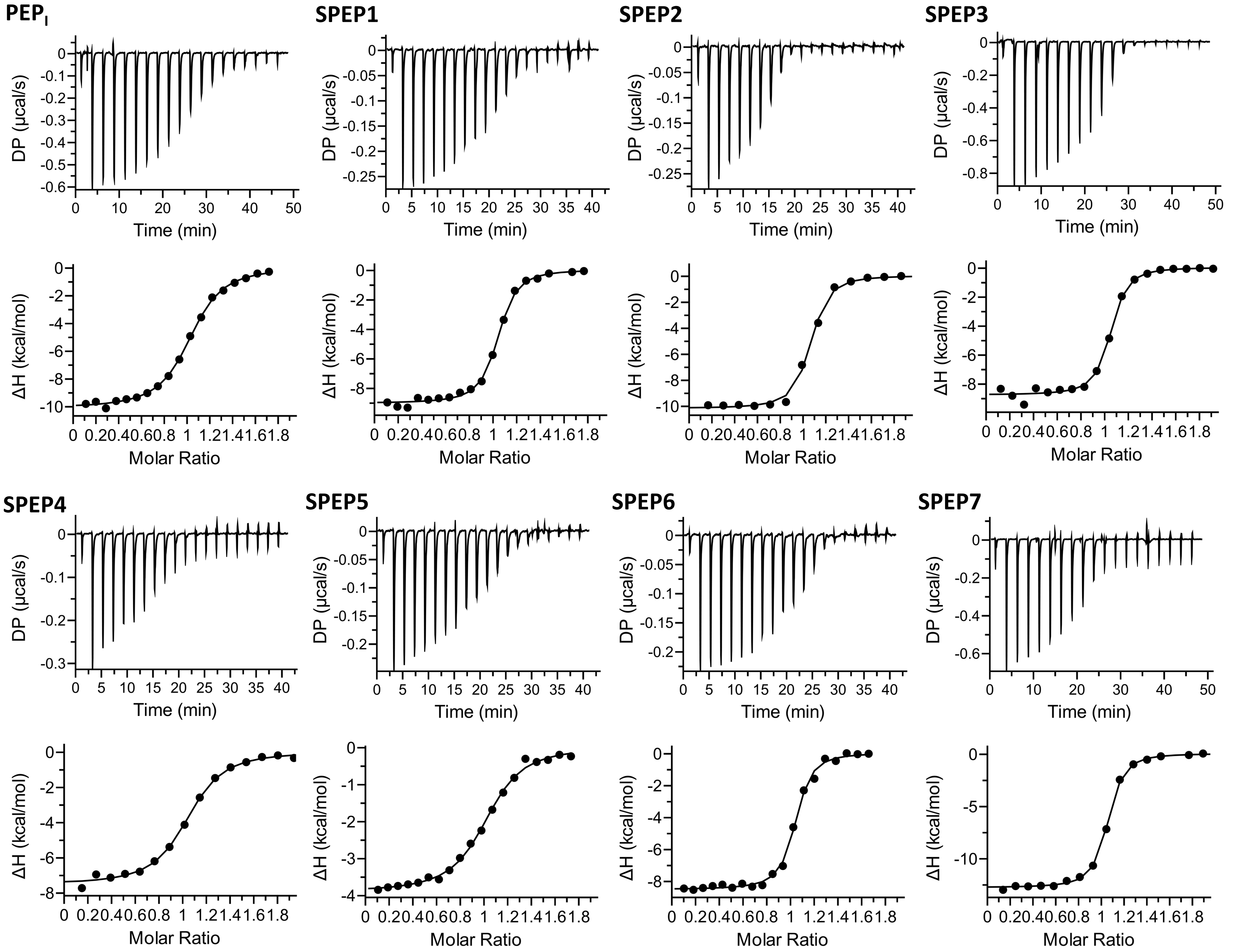


**Figure S7: ITC data for binding of $DHB_{DAXX}$ to SPEP1 to 7.**

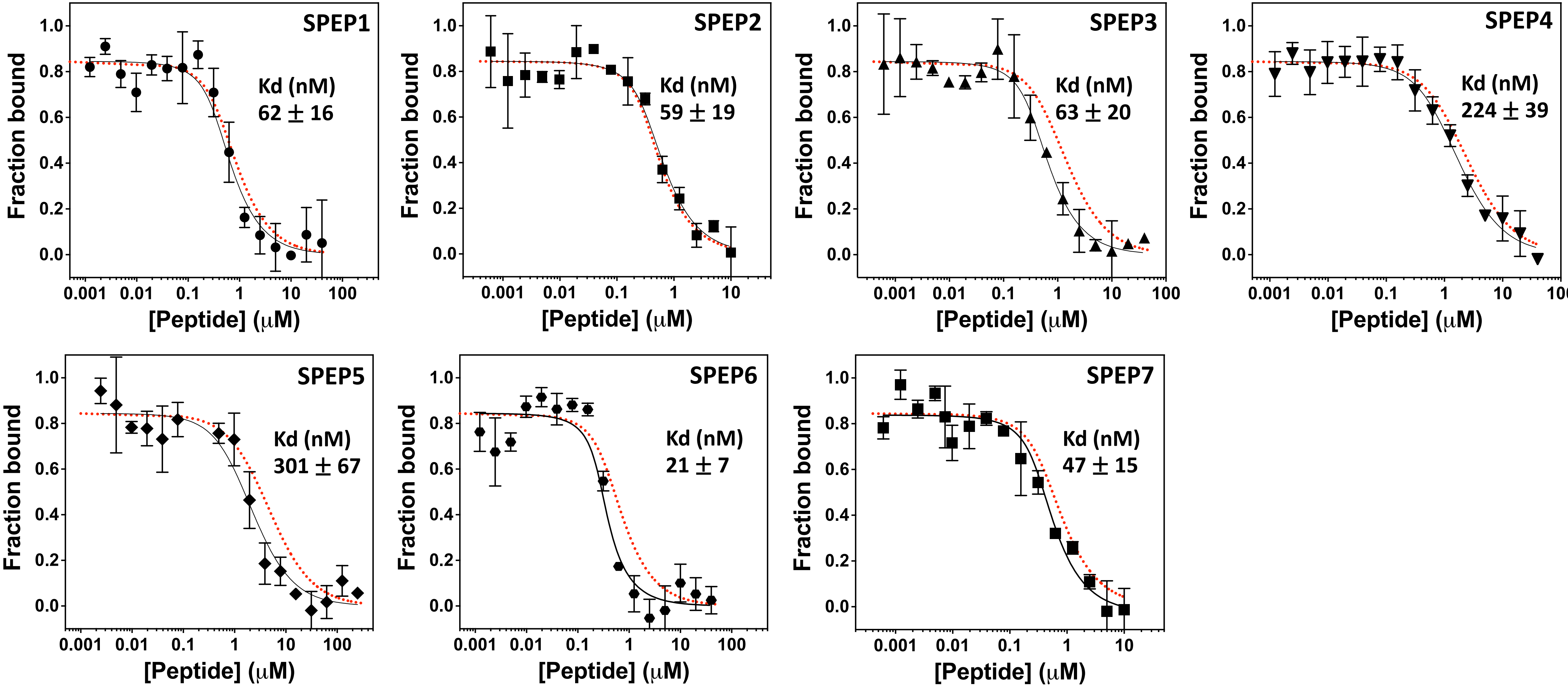


**Figure S8. Competitive displacement of FAM-SPEP7 from $DHB_{DAXX}$ by non-fluorescent SPEP peptides 1-7.**

**Figure S9. The formation of the $^{15}$N-NSIM-DHB$_{DAXX}$/FAM-SPEP7 pre-complex and subsequent binding of SUMO-1 in sequential titrations monitored using $^{1}$H-$^{15}$N HSQC NMR.**

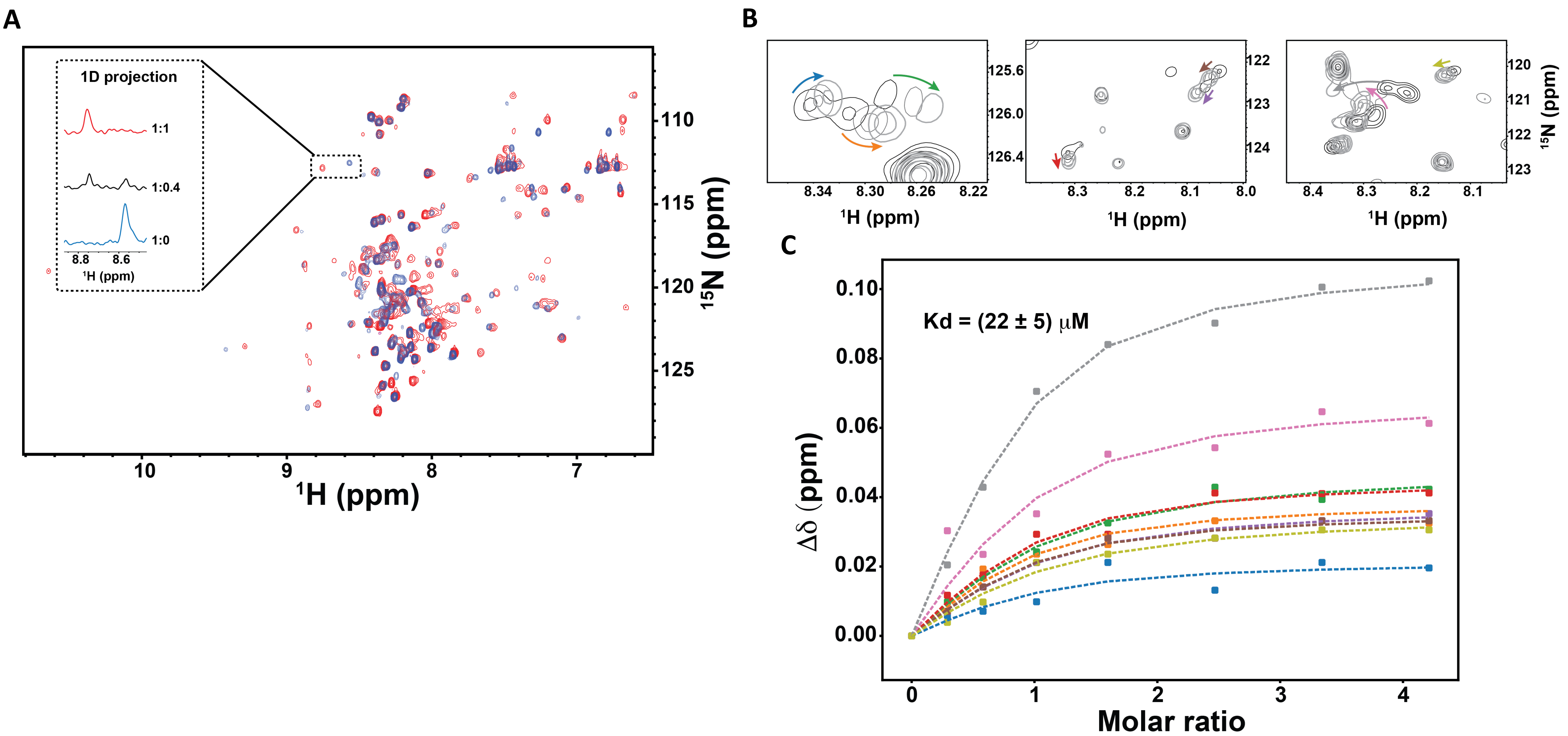

Figure S10. Cellular toxicity and localisation of stapled peptides.

A

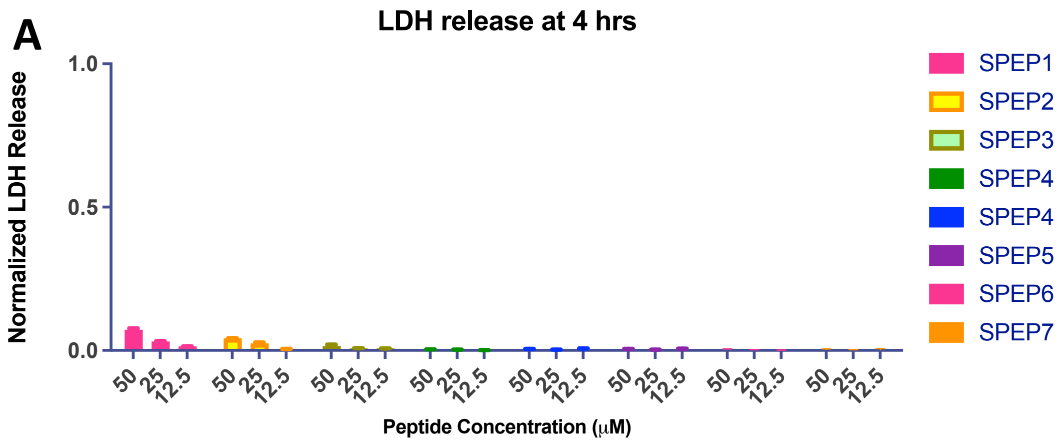


B

FAM STPEP7 24 hrs

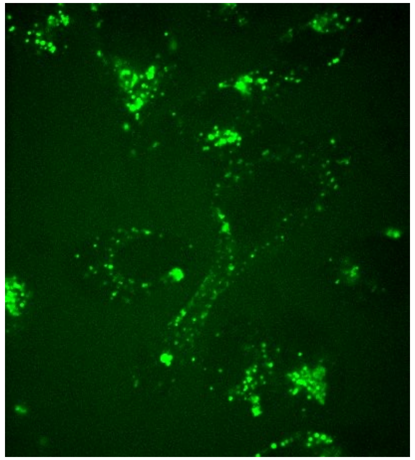

**Table S2: Binding of stitched peptides determined through computational and experimental methods. Kd values were determined from fluorescence polarization experiments.**

| **Peptide** | **Ele** | **Vdw** | **Polar** | **Nonpolar** | **ΔG*** | **Kd (nM)** |
|---|---|---|---|---|---|---|
| STPEP1 | -814.2 (52) | -63.7 (2.1) | 799.5 (47) | -11.7 (0.5) | -90.1 (7) | n.d |
| STPEP2 | -726.3 (15) | -62.1 (1.0) | 712. 0 (14) | -11.8 (0.2) | -88.2 (1.5) | 32.4 (2.6) |
| STPEP3 | -630.7 (18) | -61.0 (3) | 614.9 (17) | -11.5 (0.4) | -88.2 (4.3) | 34.9 (2.4) |
| STPEP4 | -447.7 (34) | -60.5 (1.8) | 439.9 (31) | -11.2 (0.1) | -79.5 (1.0) | 147.8 (12.4) |
| STPEP5 | -772 (36) | -55.0 (3) | 763 (32) | -10.7 (0.4) | -71.6 (7.5) | n.d |
| STPEP6 | -694 (45) | -52.2 (6) | 648.8 (44) | -10.4 (0.8) | -71.2 (9.1) | n.d |
| STPEP7 | -662.7 (51) | -57.8 (0.7) | 654.5 (45) | -10.8 (0.2) | -76.8 (5.5) | 268.9 (17.3) |

**$Binding energies are calculated using MMPBSA method. All values are calculated in kcal mol$^{-1}$ at 300K. Errors of 1 SD are shown in parentheses. *Entropy changes are not calculated, therefore the binding energy calculated here corresponds to only the enthalpy contribution.**

### Figure S11. Conformational analysis of stitched peptides.

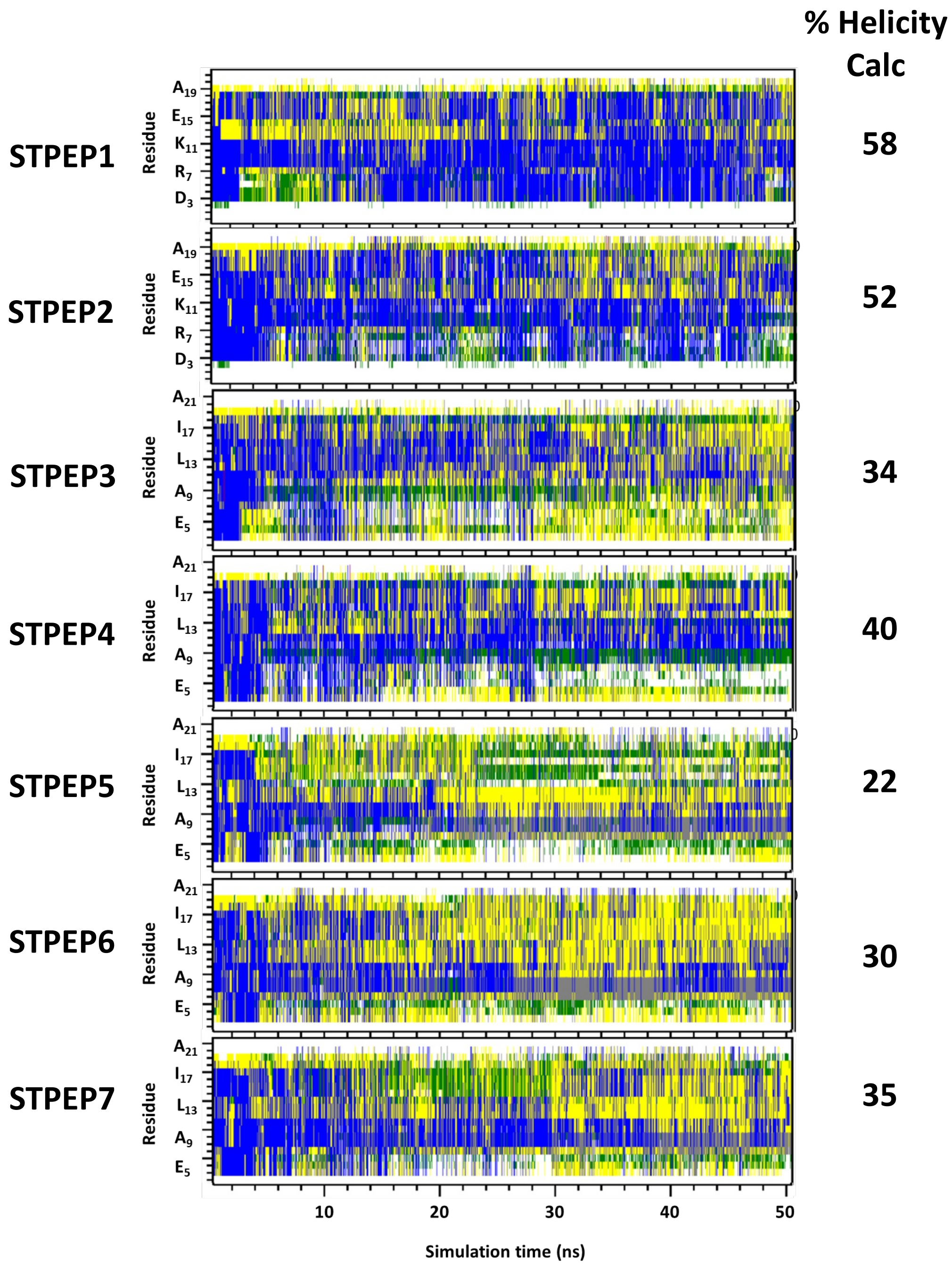

### Figure S12. Computational analysis of the stability of $DHB_{DAXX}$/STPEP complexes.

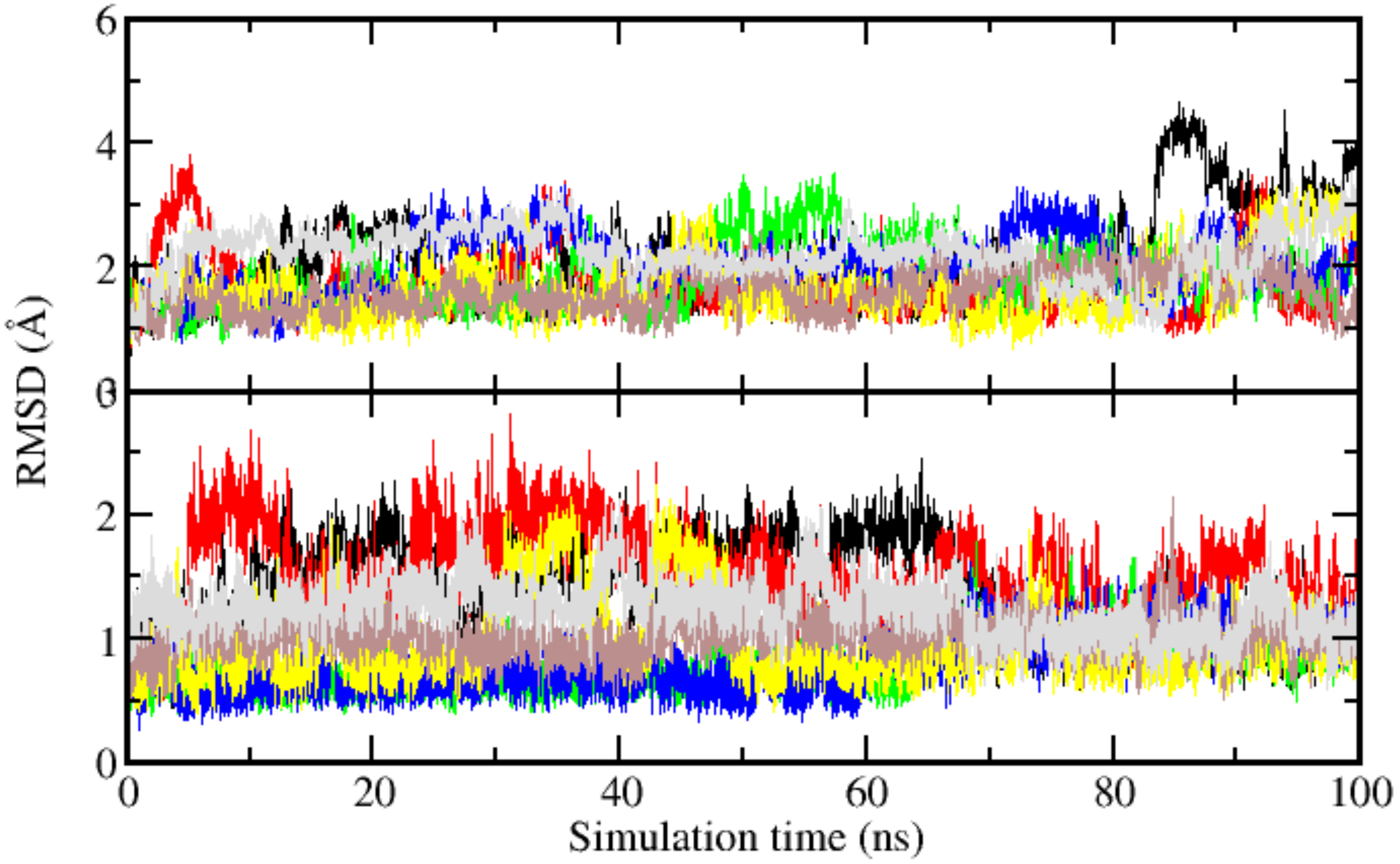
